# The Par complex regulates apical-basal cell polarity through modulation of FAK signaling homeostasis

**DOI:** 10.64898/2026.05.03.722465

**Authors:** Meiai He, Lining Liang, Yulu Wang, Yongyu Chen, Hao Sun, Lin Guo, Changpeng Li, Jingcai He, Yanhua Wu, Shiyu Chen, Tingting Yang, Fei Meng, Qiwen Ren, Linna Dong, Lin Liu, Qianqian Zou, Tianya Zhang, Xinyue Hou, Qing Guo, Dajiang Qin, Hui Zheng

**Affiliations:** Key Laboratory of Biological Targeting Diagnosis, Therapy and Rehabilitation of Guangdong Higher Education Institutes, The Fifth Affiliated Hospital of Guangzhou Medical University, Guangzhou, 510799, China; Guangdong Provincial Key Laboratory of Stem Cell and Regenerative Medicine, Guangdong-Hong Kong Joint Laboratory for Stem Cell and Regenerative Medicine, Key Laboratory of Digital Life and Medicine, Guangzhou Institutes of Biomedicine and Health, Chinese Academy of Sciences, Guangzhou, 510530, China; University of Chinese Academy of Sciences, Beijing, 100049, China; Joint School of Life Sciences, Guangzhou Medical University, Guangzhou, 511436, China; Human Cell Lineage Atlas Facility, Guangzhou Institutes of Biomedicine and Health, Chinese Academy of Sciences, Guangzhou 510530, China; Centre for Regenerative Medicine and Health, Hong Kong Institute of Science & Innovation, Chinese Academy of Sciences; Hong Kong SAR, China

**Keywords:** cell polarity, Par complex, naïve-to-primed transition, AKT–FURIN– LEFTY–FAK signaling axis, neural tube organoids

## Abstract

Cell polarity complexes are essential for embryogenesis, but their regulatory mechanisms during early developmental transitions remain incompletely understood. Here, we individually deleted the Crumbs, Par, and Scrib polarity complexes in mouse embryonic stem cells (mESCs). While loss of any single complex did not affect pluripotency or proliferation, deletion of Par complex disrupted the naïve-to-primed transition and impaired subsequent differentiation, including neural tube organoids formation. Mechanistically, Par complex deficiency led to hyperphosphorylation of focal adhesion kinase (FAK) at primed stage, driving a morphological shift from flat monolayer clusters to dome-shaped colonies. FAK inhibition rescued the aberrant morphology. Upstream, Par complex loss increased AKT phosphorylation, which acted through the FURIN–LEFTY axis, ultimately modulating FAK activity. In addition, conditioned medium from wild-type cells partially rescued differentiation defects in Par knockout cells in a LEFTY-dependent manner. These phenotypes were consistently observed in naïve-to-primed transition, neural stem cell differentiation, embryoid body formation, teratoma assays, and neural tube organoid differentiation. Together, these findings establish a Par–AKT–FURIN–LEFTY–FAK signaling cascade that links apical-basal polarity to early lineage specification and morphogenesis, providing a mechanistic framework for how polarity cues are translated into developmental outcomes.

**Graphical Abstract:** 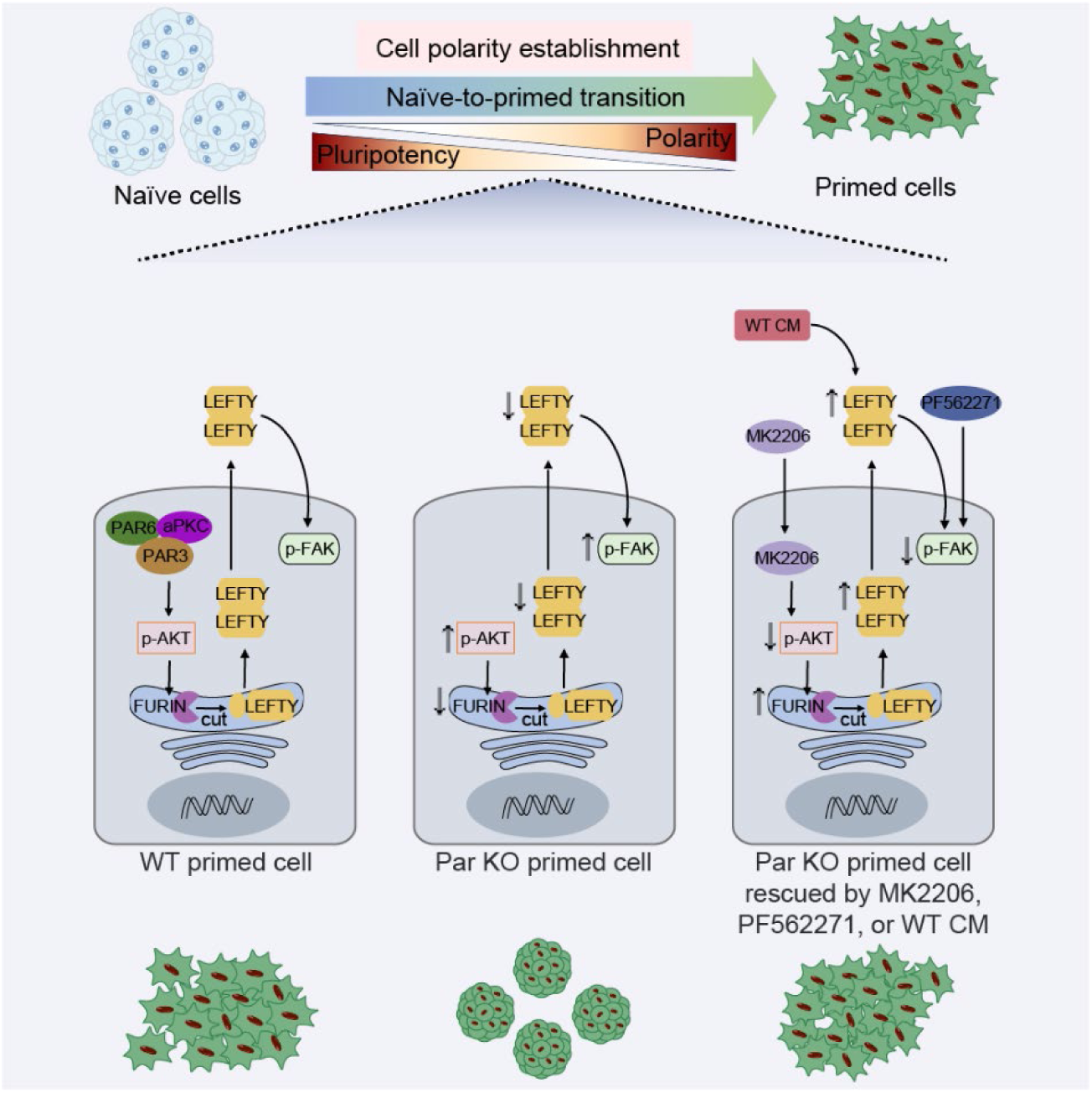

**Significance Statement:** This study elucidates the molecular mechanism by which the Par complex regulates the establishment of cell polarity. The authors demonstrate that Par complex promotes expression of the protein convertase FURIN via AKT signaling, thereby enhancing the maturation and secretion of LEFTY protein. This process ultimately regulating FAK activity and controlling the establishment of cell polarity. These findings reveal how polarity cues govern early lineage specification and morphogenesis, with implications across multiple developmental contexts.

## Introduction

The establishment of cell polarity during early embryogenesis represents a fundamental prerequisite of developmental biology, governing embryonic axis formation, cell fate determination, and tissue morphogenesis (Lamba et al., 2025; Leung et al., 2016). In mammals, apical–basal polarity is first established at the 8-cell stage, whereas in non-mammalian model organisms such as C. elegans and Drosophila, polarity is established much earlier, during oogenesis or the first cleavage divisions (Korotkevich et al., 2017; Nelson, 2003). In the mouse preimplantation embryo, the first morphological manifestation of polarity is the formation of an apical domain at the 8-cell stage, which marks the emergence of apical–basal polarity and coincides with the onset of compaction (Zhu & Zernicka-Goetz, 2020). This process involves the apical localization of conserved polarity complexes such as Crumbs and Par, while the Scrib complex becomes basolaterally enriched to maintain basal identity (Etemad-Moghadam et al., 1995; Watson et al., 2023). Apical–basal polarity is subsequently remodeled during subsequent divisions and is required for the asymmetric partitioning of fate determinants, which distinguishes the trophectoderm from the inner cell mass (Anani et al., 2014; Korotkevich et al., 2017). The Par complex—comprising Par3, Par6, and atypical protein kinase C (aPKC)—is particularly important in early mouse development, where it directs apical domain assembly and is essential for blastocyst formation and lineage specification (Plusa et al., 2005; Zhu et al., 2020). The apical domain serves as a platform for recruiting cell fate regulators and concentrating lineage-specific markers, thereby translating polarity cues into lineage-specific transcriptional programs (Zhu et al., 2021). Notably, the same polarity modules that pattern the early embryo are redeployed in pluripotent stem cell models, providing an experimentally tractable system in which to dissect how polarity contributes to potency transitions (Kalkan et al., 2017).

The naïve-to-primed transition (NPT) represents a critical developmental phase in pluripotent stem cells, marked by distinct morphological and molecular changes. Naïve cells maintain dome-shaped colonies and express high levels of pluripotency factors such as *Klf4* and *Esrrb*, whereas primed cells adopt flat monolayer morphology and upregulate lineage-commitment genes including *Brachyury* (*T*) and *Fgf5* (Tesar et al., 2007; Weinberger et al., 2016). Developmentally, naïve state mirrors the pre-implantation inner cell mass, while primed state resembles the post-implantation epiblast (Kalkan et al., 2017). Although naïve cells possess broad differentiation potential, they must transition through the primed state to respond to differentiation signals (Hayashi & Saitou, 2013; Rostovskaya et al., 2019). Primed cells thus represent an intermediate stage between pluripotency and lineage commitment, exhibiting enhanced lineage responsiveness and cell polarity (Brons et al., 2007). How this polarity is acquired, and whether it instructs or merely accompanies the NPT, remains an open question central to the present work.

Early studies identified key roles for pathways such as Hippo and AKT in polarity regulation (Tokamov et al., 2023; Xu et al., 2019). More recent interdisciplinary work integrating biomechanical and biomolecular analyses has revealed that extracellular matrix (ECM) and phase separation of polarity proteins collaboratively regulate cell adhesion and migration (Dias Gomes & Iden, 2021; Guzman-Herrera & Mao, 2020). During migration, focal adhesion kinase (FAK) exhibits spatially polarized activity within focal adhesions (Li et al., 2023). FAK amplifies its own activation and membrane localization in response to mechanical or integrin-mediated signals, triggering downstream effectors to promote cell polarity (Diaz-Palacios et al., 2025; Y. Wang et al., 2024). Additionally, active FAK recruits the PI3K–AKT signaling complex to establish a polarity axis and sustain migratory directionality (J. Wang et al., 2024).

Loss of cell polarity is implicated in various human diseases, including neurological disorders, cancer, and tissue fibrosis (Zhan et al., 2008; Zhang & Wei, 2022), underscoring the need to define how polarity is established and maintained in pluripotent cells.

In this study, we used CRISPR-Cas9 to knock out Crumbs, Par, and Scrib polarity complexes in mouse embryonic stem cells (ESCs). We systematically characterized polarity abnormalities in multiple differentiation systems, and combined pharmacological inhibition, co-culture and conditioned-medium rescue approaches to test how polarity disruption impairs the naïve-to-primed transition and downstream differentiation. Our findings uncover key molecular mechanisms through which cell polarity is established and controls the potency transition of pluripotent stem cells.

## Results

### The Par complex is required for morphological remodeling during NPT

To investigate the roles of polarity complexes in cell polarity establishment, we used CRISPR-Cas9 to knock out Crumbs, Par, or Scrib complex in OG2 embryonic stem cells (OG2-ESCs), which represent naïve cells (Fig. 1A and S1A). Two independent knockout cell lines per complex were established and validated for subsequent experiments (Fig. S1B).

**Figure 1.**
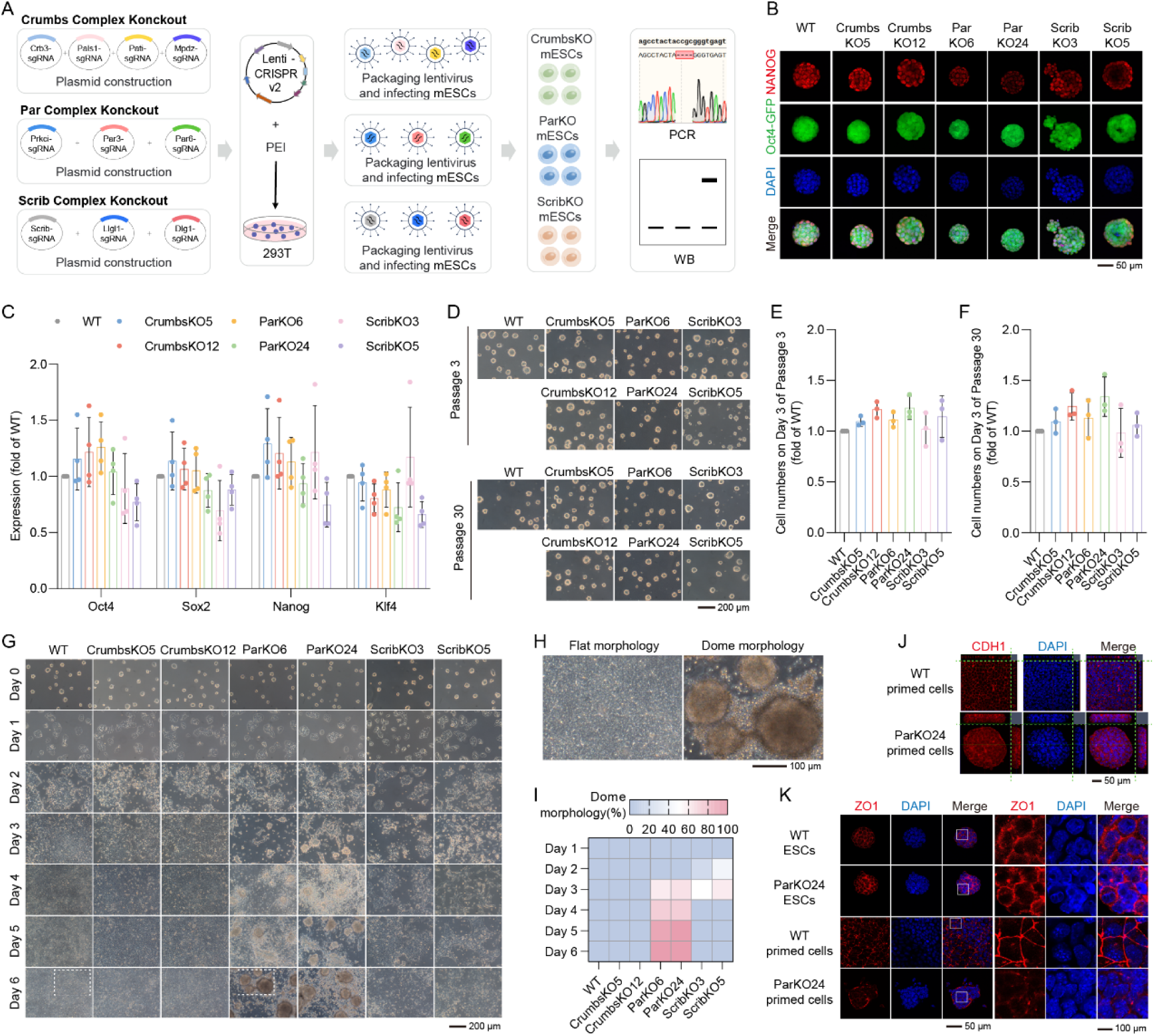
Par complex is essential for NPT. **(A)** Schematic illustration of the generation of ESCs without Crumbs, Par, or Scrib complex. **(B-F)** Crumbs, Par, or Scrib complex KO did not affect ESCs pluripotency, morphology, or proliferation. The expression levels of endogenous pluripotency markers *Nanog*, *Oct4*, *Sox2*, and *Klf4* showed no significant differences **(B-C)**. ESCs maintained a characteristic dome-shaped colony morphology over more than 30 passages **(D)**. The initial cell numbers were 35,000 on day 0, and cell numbers were determined on day 3 **(E-F)**. **(G-I)** Par KO yielded cells exhibiting a dome-shaped colony morphology during NPT, contrasting with the flat monolayer morphology of WT cells. The defining criteria for both morphological types are specified in **(H)**, and the proportion of dome-shaped colony morphology in each group of cells is shown in **(I)**. **(J-K)** Par KO resulted in aberrant subcellular localization of CDH1 and ZO1 in primed cells, as captured by LSM800 confocal microscopy. Experiments were repeated at least three times unless otherwise stated. Error bars represent S.D. One-way ANOVA was performed in **(C)**, **(E)**, and **(F)**; no significant differences were observed among any of the group comparisons.

We first assessed whether loss of polarity complexes affects pluripotency. Immunofluorescence analysis showed that knockout ESCs maintained normal expression of the pluripotency markers, and 99% of them were Oct4-GFP^+^ (Fig. 1B and S1C). RT-qPCR further confirmed that mRNA levels of *Oct4*, *Sox2*, *Nanog*, and *Klf4* were not significantly altered in any of the knockout cell lines (Fig. 1C). Moreover, all knockout ESCs retained normal morphology and proliferation capacity after at least 30 passages (Fig. 1D and 1F). These results indicate that deletion of polarity complexes does not affect pluripotency maintenance of ESCs.

During NPT process, Par complex knockout cells (ParKO6 and ParKO24) displayed morphology similar to WT cells during the first two days, appearing as a flat monolayer morphology (Fig. 1G). By day 3, however, Par knockout (Par KO) cells began to disperse and form dome-shaped colony, a phenotype that became more pronounced over time. By day 6, over 90% of Par KO cells exhibited dome-shaped colonies (Fig. 1G and 1I). These dome-shaped colonies persisted even after three passages (Fig. S1D and S1E), indicating a failure to establish epithelial monolayer organization. Live-cell imaging revealed that from day 3 onward, Par KO cells tended to migrate and aggregate into dome-shaped colonies, suggesting day 3 is a crucial time point for the transformation of cell morphology (Vid. 1 and 2).

In contrast, Scrib complex knockout cells (ScribKO3 and ScribKO5) showed approximately 10% dome-shaped colonies on day 3 of NPT but reverted to flat monolayer clusters by day 6 (Fig. 1G and 1I). Crumbs complex knockout cells (CrumbsKO5 and CrumbsKO12) displayed no notable morphological differences compared to WT throughout NPT (Fig. 1G and 1I). These results establish the Par complex as essential for morphological establishment during NPT. Given the consistent phenotypes in ParKO6 and ParKO24, we focused subsequent studies on ParKO24.

Cell junctions represent hallmark structural features of epithelial polarity, with adherens junctions (marked by CDH1) and tight junctions (marked by ZO1) requiring proper apical polarity for their assembly and function in regulating cell fate (Fleming et al., 1989). To assess junctional organization in our model, we performed immunofluorescence analysis of CDH1 and ZO1 in Par KO primed cells. On day 6 of NPT, CDH1 was localized at the cell-cell junctional membrane in WT primed cells, consistent with its characteristic apical distribution at tight junctions (Yonemura et al., 1995), whereas in Par KO primed cells, CDH1 was extensively distributed around the whole cell membrane without distinct junctional enrichment (Fig. 1J). Additionally, ZO1 showed no significant difference between Par KO and WT ESCs, but its distribution pattern differed significantly in primed cells: WT cells showed strong and continuous ZO1 signal at the apical membrane domain, while Par KO primed cells exhibited weak and scattered staining distributed along the plasma membrane (Fig. 1K). These findings demonstrate that the Par complex is essential for proper junctional organization and polarity establishment during NPT.

### The Par complex regulates morphological remodeling through the AKT and FAK signaling pathways

To elucidate the molecular mechanisms by which Par complex regulates cell polarity establishment, we performed RNA-seq on WT and Par KO cells at five time points (days 0, 1, 3, 4, and 6) during NPT (Fig. 2A).

**Figure 2.**
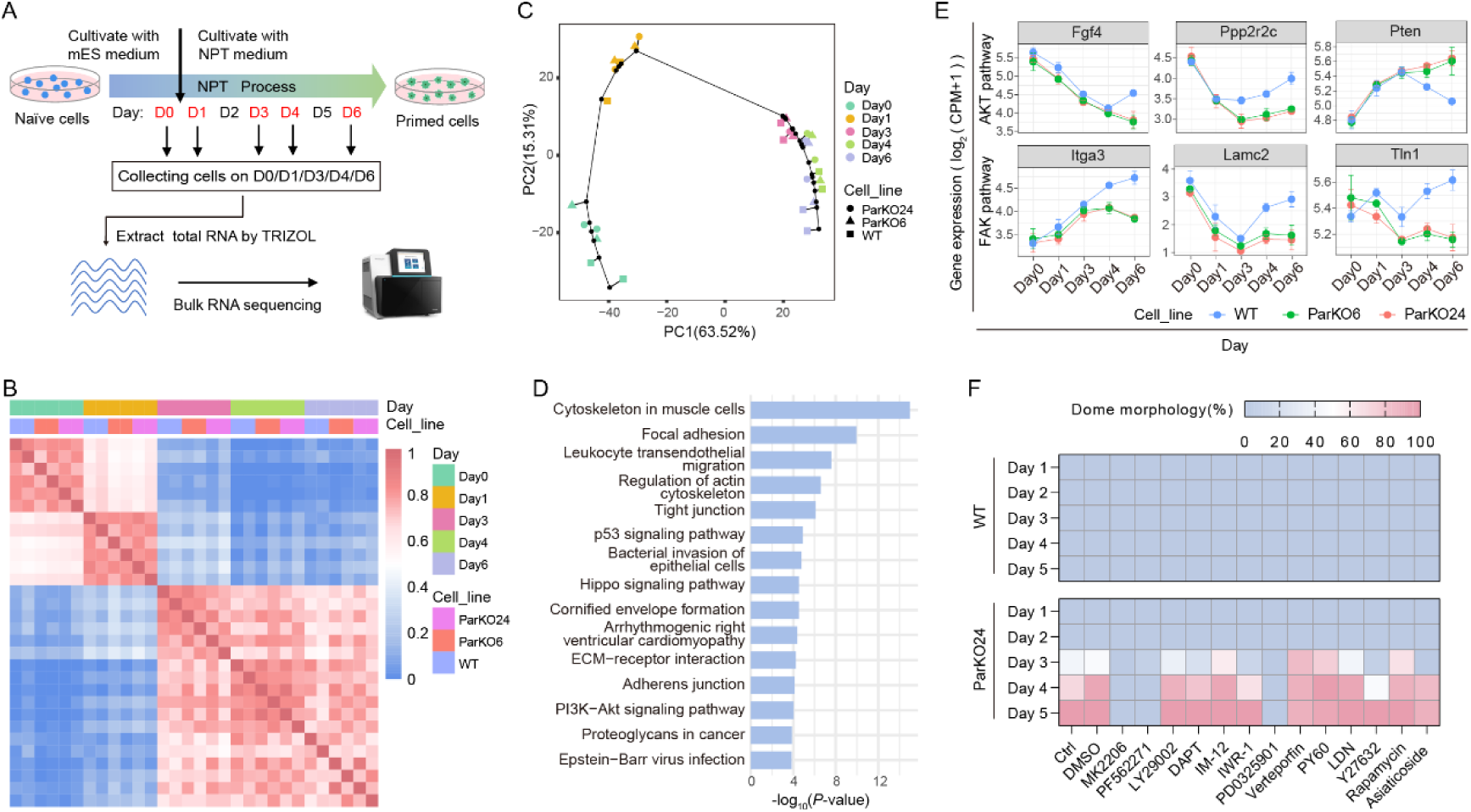
Suppression of AKT or FAK signaling rescues the defect induced by Par KO. **(A)** Schematic illustration of RNA-seq analysis. Cells on day 0, 1, 3, 4, and 6 during NPT were collected for bulk RNA-seq. **(B)** Heatmap analysis of sample-to-sample correlations of the transcriptomics of Par KO and WT cells during NPT. **(C)** Pseudotime analysis of transcriptomics of WT and Par KO cells during NPT. **(D)** KEGG analysis of enriched pathways of DEGs on day 6. **(E)** Expression of key genes of the enriched pathways in (D) was analyzed, also see Figure S2B. **(F)** Cells were treated with agonists or inhibitors targeting the enriched pathways during NPT, followed by morphological analysis of dome-shaped colonies. Experiments were repeated at least three times, with the exception of high-throughput sequencing, or unless otherwise stated. Error bars represent S.D.

Heatmap and pseudotime analysis indicated that major transcriptional shifts occurred at days 1 and 3 in both genotypes (Fig. 2B and 2C). The overall differentiation trajectory of Par KO cells resembled that of WT, consistent with their ability to upregulate primed markers (Fig. 2C). RNA-seq analysis revealed that the number of differentially expressed genes (DEGs) between Par KO and WT cells increased progressively from day 3 onward, with 160 downregulated and 363 upregulated genes at day 3, and 945 downregulated and 948 upregulated genes at day 6 in Par KO cells (Fig. S2A). KEGG analysis revealed that downregulated genes were enriched in focal adhesion, Hippo signaling, PI3K-AKT signaling, cytoskeleton regulation, cell junction, and ECM-receptor interaction (Fig. 2D). Key genes in these pathways, including AKT and FAK, showed differential expression as early as day 3, and the differences gradually increased as NPT proceeded (Fig. 2E and S2B).

Given the broad signaling alterations, we screened small-molecule modulators of these pathways during NPT. The concentrations of small-molecule modulators were optimized based on the recommended IC_50_/EC_50_ for cell experiments to ensure no significant effects on cell proliferation or apoptosis. Among the compounds tested, only MK2206 (AKT inhibitor) and PF562271 (FAK inhibitor) rescued the morphology defects in Par KO primed cells, restoring flat monolayer clusters (Fig. 2F and S2C). No rescue was observed with compounds targeting PI3K, mTOR, Rho/ROCK, Wnt/β-catenin, Notch, BMP, TGF-β, or Hippo signaling.

Notably, Par KO primed cells treated with PD0325901 (MEK inhibitor) displayed a morphology distinct from both the flat monolayer clusters of typical primed cells and the dome-shaped colonies of naïve cells. Instead, they formed multi-layer clusters characterized by a central dome-shaped region surrounded by peripheral flat cells, exhibiting a more compact and aggregated architecture (Fig. S2C). Further detection revealed that Par KO primed cells treated with PD0325901 retained high percentage of Oct4-GFP^+^ cells (Fig. S2D) and expressed high levels of *Esrrb* and *Klf4*, and low levels of *Fgf5* (Fig. S2E). Consistent with prior reports that PD0325901 sustains pluripotency and prevents ESC differentiation (Ying et al., 2008), these results suggest that the observed “rescue” by PD0325901reflects only a morphological improvement, with maintenance of the ESC state, thereby hindering NPT progression.

In contrast, treatment with MK2206 or PF562271 fully restored the dome-shaped morphology of Par KO cells to WT-like, without significantly altering naïve and primed gene expression. This was evidenced by comparable levels of primed markers, suppression of ESC markers, and downregulation of endogenous Oct4-GFP (Fig. S2D and S4F). Together, these results indicate that inhibition of AKT or FAK signaling specifically rescued morphological defects resulting from Par complex loss during NPT.

### Conditioned medium from WT cells rescues the defects induced by Par KO

While the spatiotemporal dynamics of polarity complexes play a pivotal role in directing cell fate specification and lineage differentiation, how their mislocalization or dysregulation in individual cells affects cell fate and behavior remains unclear. To address this, we performed NPT in co-culture systems containing both WT and Par KO ESCs. When cultured alone, Par KO primed cells consistently exhibited dome-shaped colonies. In contrast, co-culture with WT ESCs at initial ratios of 1:1 or 2:1 (Par KO:WT), over 90% cells exhibited flat monolayer clusters (Fig. S3A and S3B). This rescue could be attributed to either the exclusion or the active correction of Par KO cells by WT cells.

To distinguish these possibilities, we established tdTomato-labeled Par KO ESCs and co-cultured them with WT ESCs. When cultured alone, over 92% of Par KO primed cells were tdTomato⁺ (Fig. 3A and 3B), but they proliferated more slowly than WT primed cells, because the cell numbers of Par KO primed cells were approximately 81.6% of those of WT (Fig. 3C). At the end of NPT, approximately 24.3% and 41.9% of the cells were tdTomato⁺ when Par KO and WT ESCs were co-cultured at ratios of 1:1 and 2:1, respectively (Fig. 3B), marking actual percentage of Par KO primed cells was 26.4% and 45.6% (Fig. 3D), which suggested significant survival of Par KO cells with approximately 58.7% and 73.4%. Thus, high percentage of flat morphology was possibly due to the active correction of the Par KO phenotype by WT cells.

**Figure 3.**
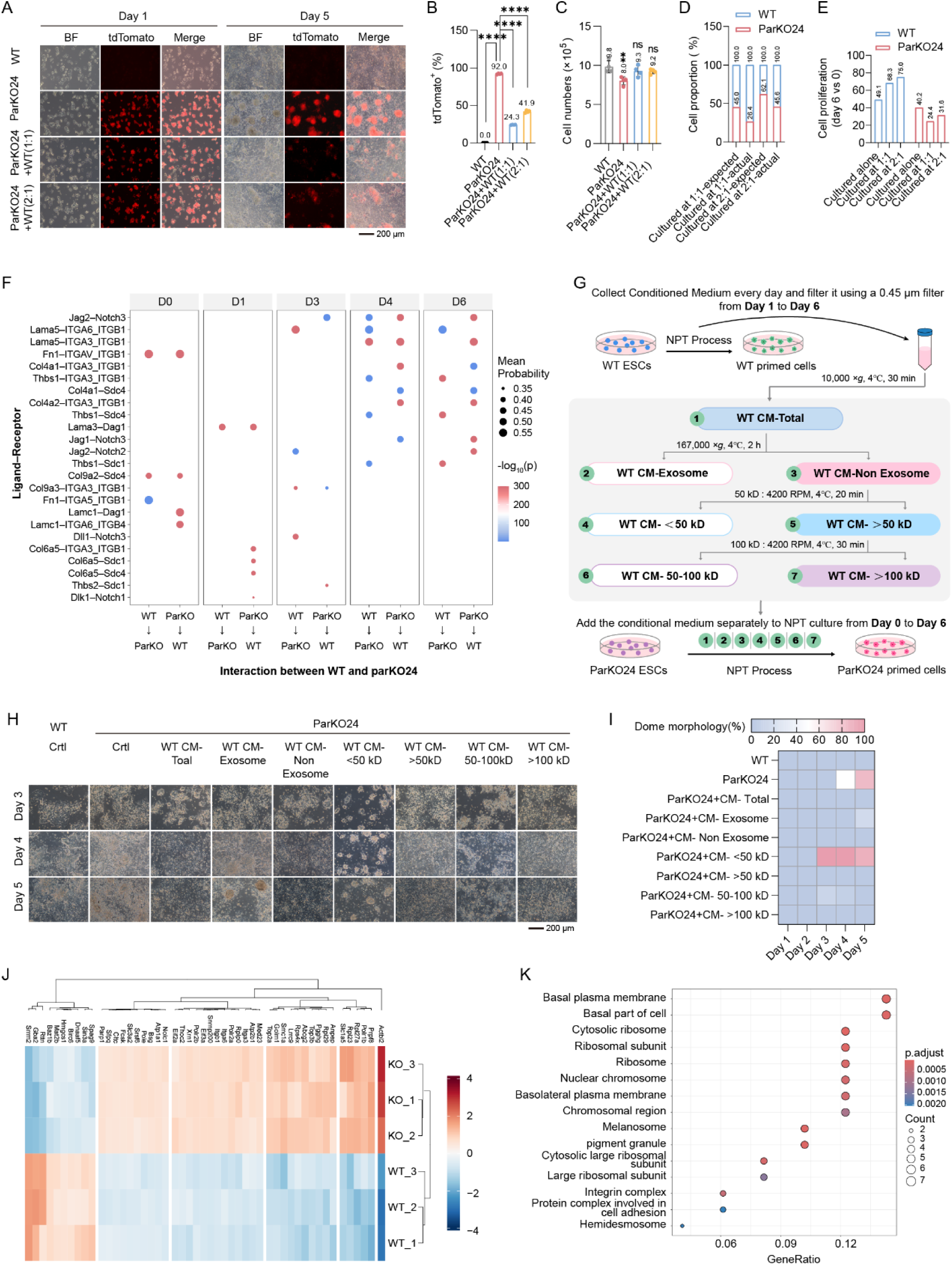
Conditioned medium from WT cells rescues the defect induced by Par KO. **(A-C)** tdTomato-labeled Par KO ESCs were co-cultured with WT ESCs during NPT (cell numbers were 20,000 on day 0). Cell numbers and the proportion of tdTomato^+^ were determined on Day 6. **(D-E)** Based on the results from (B and C), the proportion of WT and Par KO cells in the co-culture systems was calculated, along with the proliferation efficiency of WT and Par KO cells. **(F)** Analysis of interactions between Par KO and WT cells using RNA-seq data. **(G)** Schematic illustration of component separation of WT CM. The total WT CM was separated into 7 groups. **(H-I)** Different components of WT CM were used to treat the Par KO cells respectively, and the cell morphology were analyzed. **(J-K)** Protein profile was conducted on WT and Par KO CM to analyze the differentially expressed proteins and their signaling pathways. Experiments were repeated at least three times unless otherwise stated. Error bars represent S.D. One-way ANOVA was performed in **(B)** and **(C)**.

On the other hand, the proliferation of Par KO cells was impaired in co-culture systems. Given the proliferation rates of Par KO and WT cells when cultured alone, the expected percentages of Par KO cells are approximately 45.0% and 62.1% in the 1:1 and 2:1 ratio systems, respectively (Fig. 3D). Reductions around 41.3% (1:1) and 26.6% (2:1) suggested that although WT cells rescued the morphology of Par KO cells, they also competed with and impaired the growth of Par KO cells. Compared to the proliferation rate when culture alone (approximately 49-fold), the proliferation rate of WT cells in co-culture systems increased to approximately 68-fold (1:1) and 75-fold (2:1), whereas the proliferation rate of Par KO cells decreased from 40-fold in culture alone to 24-fold (1:1) and 32-fold (2:1) in co-culture systems (Fig. 3E).

These findings suggest that, at the cellular level, normal cells can either rescue the defective phenotypes of abnormal cells or increase their own population proportion to dilute the relative abundance of impaired cells. Analysis of cell-cell interactions during NPT in WT and Par KO cells revealed that the predominant interactions were associated with LAMININ and COLLAGEN pathways, exemplified by receptor-ligand pairs such as *Lama5*–*Itga6*/*Itgb1*, *Lama5*–*Itga3*/*Itgb1*, and *Col4a1*–*Itga3*/*Itgb3* (Fig. 3F). These interactions were notably strengthened between day 4 and day 6 and suggested that Par complex regulates ECM assembly and provides mechanical cues along with integrin-mediated signaling, thereby promoting cell polarization and morphology.

We next asked whether the abilities of WT cells to rescue the morphology of Par KO cells requires direct contact or is mediated by secreted factors. Conditioned medium (CM) from WT cells during days 1–6 of NPT—but not from Par KO cells or WT primed cell lysates (CL)—fully restored flat monolayer morphology in Par KO primed cells (Fig. S3C and S3D). This effect was dependent on sustained exposure; WT CM had to be applied at least from day 1 to day 4 of NPT to achieve complete rescue, whereas WT CM collected at day 0 of NPT showed no rescuing activity (Fig. S3E).

To identify the key components in WT CM, we fractionated WT CM into exosome and soluble content, and the soluble content was further separated by molecular weight (Fig. 3G). All the fractions were used to treat Par KO cells during NPT. The rescue activity was retained in groups of WT CM-Total, WT CM-Non exosome, WT CM-> 50 kDa, and WT CM-> 100 kDa (Fig. 3H and 3I). Although the 50–100 kDa fraction also eventually rescued the phenotype, its effect was delayed, indicating that the key factor(s) reside in the > 100 kDa fraction. Furthermore, the WT CM-Exosome exhibited limited rescue efficacy, particularly during the later stages of NPT. Comparative proteomic profiling of WT and Par KO CM (the > 100 kDa fractions) revealed that WT CM was enriched in proteins involved in cell organization, integrin, and cell adhesion (Fig. 3J and 3K). These findings suggest that WT cells secrete high-molecular-weight proteins or complexes that restore polarity and morphology in Par KO cells.

### The Par complex regulates morphological remodeling through an AKT–FURIN– LEFTY–FAK signaling axis

To determine whether AKT inhibition and WT CM share a common rescue mechanism, we performed RNA-seq on Par KO primed cells treated with either MK2206 or WT CM at day 6 of NPT. Clustering analysis demonstrated that both treatments shifted the transcriptional profiles of Par KO primed cells toward the WT pattern, showing clear separation from untreated Par KO controls (Fig. 4A). Compared to Par KO primed cells, 787, 509 and 585 genes were upregulated in WT, MK2206-and WT CM-treated cells respectively. Among these, 302 genes were commonly upregulated under both rescue conditions, indicating substantial mechanistic overlap (Fig. 4B). A core set of 198 genes was upregulated across all three groups (WT, MK2206, and WT CM) compared to Par KO. KEGG analysis revealed that these 198 genes were primarily enriched in pathways related to cytoskeleton, focal adhesion, and ECM-receptor interaction (Fig. 4C), suggesting that both MK2206 and WT CM rescue morphological defects by regulating cell adhesion and ECM-related processes.

**Figure 4.**
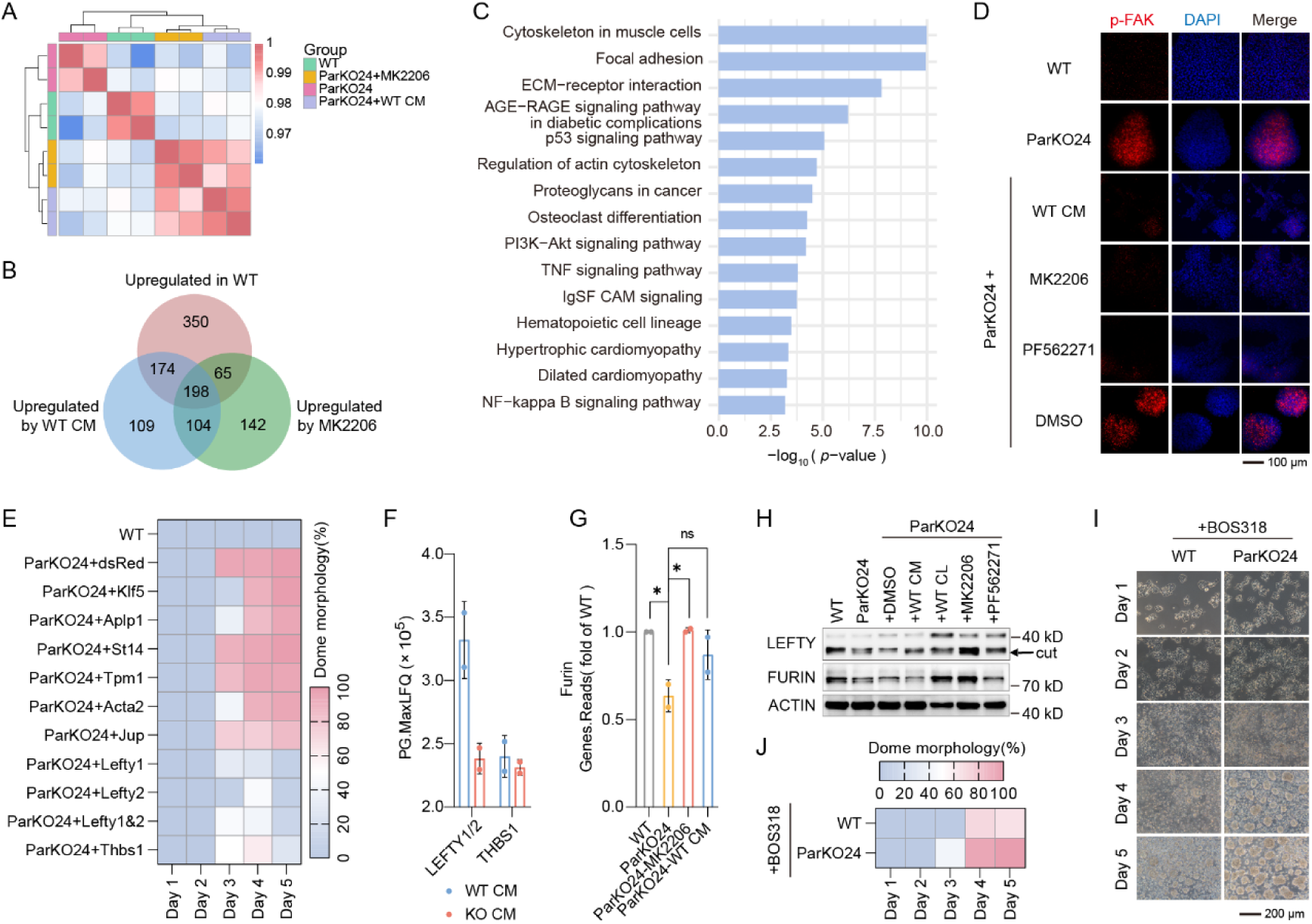
The AKT–FURIN–LEFTY–FAK signaling axis mediates Par complex-dependent morphological remodeling. **(A)** Transcriptome patterns of Par KO primed cells treated with MK2206 and WT CM were analyzed. **(B)** Overlap analysis was performed on the upregulated genes from three comparisons: WT versus Par KO, MK2206-or WT CM-treated Par KO versus untreated Par KO. **(C)** KEGG analysis was then conducted on the 198 commonly upregulated genes in **(B)**. **(D)** Par KO ESCs were treated with WT CM, MK2206, or PF562271 respectively during NPT, and p-FAK levels were detected on Day 6. **(E)** Target proteins were overexpressed individually in Par KO ESCs, followed by subjecting the overexpressing ESCs to NPT and analyzing their cell morphology. **(F)** The levels of LEFTY1/2 and THBS1 in WT and Par KO CM were analyzed. **(G)** Par KO ESCs were treated with MK2206 or WT CM respectively during NPT, and the *Furin* mRNA expression was detected on Day 6. **(H)** Par KO ESCs were treated with WT CM, MK2206, or PF562271 respectively, and the protein level of the LEFTY and FURIN were detected. The position indicated by the arrow represents the LEFTY cut by FURIN. **(I-J)** Cell morphology of WT and Par KO cells treated with *Furin* inhibitor (BOS318) during NPT. Experiments were repeated at least three times unless otherwise stated. Error bars represent S.D. One-way ANOVA was performed in **(G)**.

Considering that MK2206 is an AKT inhibitor, we measured phosphorylated AKT (p-AKT) levels across different groups. Par KO led to upregulated p-AKT levels, and this increase was effectively suppressed by MK2206 treatment (Fig. S4A). Notably, treatment with WT CM did not decrease p-AKT levels, suggesting WT CM might perform rescue function downstream of AKT signaling. The FAK signaling pathway serves as a central node that integrates upstream inputs from both PKC and AKT pathways, while transducing extracellular cues derived from ECM-integrin interactions into intracellular signaling cascades (Sakthivel et al., 2026). Its enrichment here suggests a central role in rescuing Par KO cell morphology (Fig. 4C). Consistent with this, expression of the focal adhesion proteins *Actn1* and *Flnb*, the integrin *Itga5*, and the ECM regulator *Thbs1* was significantly reduced in Par KO primed cells at day 6 of NPT (Fig. S4B). Importantly, FAKwas hyperphosphorylated in Par KO primed cells compared to WT primed cells, yet both MK2206 and WT CM effectively reduced phosphorylated FAK (p-FAK) levels (Fig. 4D).

The FAK inhibitor PF562271, which significantly rescued the NPT defects caused by Par KO, effectively reduced p-FAK levels in Par KO primed cells (Fig. 4D). Given that both MK2206 and WT CM also attenuated the elevated p-FAK, we propose that all three treatments restore the flat monolayer morphology in these cells by regulating FAK signaling homeostasis. To validate this signaling pathway, we performed experiments using an additional Par KO clone, ParKO6. Similarly, WT CM, MK2206 and PF562271 treatment each rescued the NPT defects in ParKO6 (Fig. S4C and S4D), and all three treatments downregulated the hyperactivated p-FAK (Fig. S4E). Moreover, cells under all three treatments underwent a normal NPT process, characterized by low expression of naive markers and high expression of primed markers (Fig. S4F).

Through integrated analysis of proteomic and RNA-seq data, we identified a set of functional proteins associated with the FAK signaling (Fig. S4G). These candidate proteins were subsequently overexpressed in Par KO ESCs. Overexpression of LEFTY1, LEFTY2, or THBS1 in Par KO ESCs significantly rescued cell morphology during NPT, with LEFTY1 and LEFTY2 showing the strongest effects (Fig. 4E and S4H). Actually, the protein levels of LEFTY were significantly higher in WT CM compared to KO CM (Fig. 4F), suggesting that WT cells modulate FAK signaling via secretion of LEFTY proteins. Under these conditions, WT CM treatment supplied extracellular LEFTY to Par KO ESCs, thereby rescuing the phenotypic defects of Par KO primed cells. It is therefore reasonable to infer that MK2206 rescues the defects in Par KO primed cells through upregulation of LEFTY expression, which in turn suppresses p-FAK. Concordantly, LEFTY overexpression reduced p-FAK in Par KO primed cells (Fig. S4I), consistent with reported roles of LEFTY in suppressing FAK signaling (Alowayed et al., 2016). These results suggest that MK2206 and WT CM rescue morphological defects by suppressing excessive p-FAK through LEFTY.

The intracellular LEFTY proprotein requires proteolytic cleavage by FURIN in the Golgi apparatus to be secreted extracellularly as a stable homodimer, which then exerts its biological function (Dubois et al., 2001). To investigate whether MK2206 influences FURIN expression, we analyzed transcriptomic data and found that Par KO downregulated *Furin* mRNA expression, whereas MK2206 treatment restored its transcript levels in Par KO cells (Fig. 4G). Consistent with this, MK2206 also increased FURIN protein abundance, consequently enhancing the production of cut LEFTY protein (Fig. 4H and S4J). To further validate the essential role of FURIN-mediated cleavage of LEFTY, we treated both WT and Par KO ESCs with BOS318, a highly specific and potent inhibitor of FURIN that irreversibly binds to the protease by mimicking its natural substrate (Ivachtchenko et al., 2024). BOS318 treatment induced WT primed cells to adopt a dome-shaped morphology resembling that of Par KO primed cells (Fig. 4I and 4J), and cleaved LEFTY protein was markedly reduced (Fig. S4K). These observations demonstrate that inhibition of FURIN proteolytic activity by BOS318 prevents the secretion and functional maturation of LEFTY, thereby abrogating its ability to suppress p-FAK and leading to aberrant primed cell morphology. Following its secretion into the extracellular space, how does LEFTY regulate intracellular p-FAK homeostasis? Integrated analysis of both transcriptomic and proteomic data revealed that Par KO leads to significant enrichment of pathways associated with ECM and integrin (Fig. 2D, 3F, 3K, and S4B). Notably, both MK2206 and WT CM treatment co-upregulated the ECM-receptor interaction pathway (Fig. 4C). We therefore propose that secreted LEFTY acts as an extracellular signal that activates specific ECM receptors and modulates integrin complexes. This in turn regulates FAK phosphorylation, thereby maintaining normal cell adhesion and morphology. To test this hypothesis, we subjected LEFTY-overexpressing Par KO ESCs to NPT, followed by qPCR analysis. During NPT, Par KO led to downregulation of the mRNA levels of ECM- and integrin-related components, whereas LEFTY overexpression significantly upregulated the mRNA levels of these components (Fig. S4L).

These findings collectively demonstrate that Par complex suppresses p-AKT to upregulate FURIN expression. Following FURIN-mediated proteolytic maturation, LEFTY is secreted and ultimately contributes to FAK signaling homeostasis, thereby promoting cell polarity establishment.

### The Par complex is required for differentiation across multiple developmental contexts via the AKT–FAK signaling axis

Given that Par complex loss disrupts cell morphology during NPT, we asked whether it also impairs multi-lineage differentiation. We differentiated WT ESCs, Par KO ESCs, WT primed cells, and Par KO primed cells into embryoid bodies (EBs) using suspension culture. On day 6, EBs derived from four types of cells formed spherical structures with comparable morphological characteristics (Fig. 5A).

**Figure 5.**
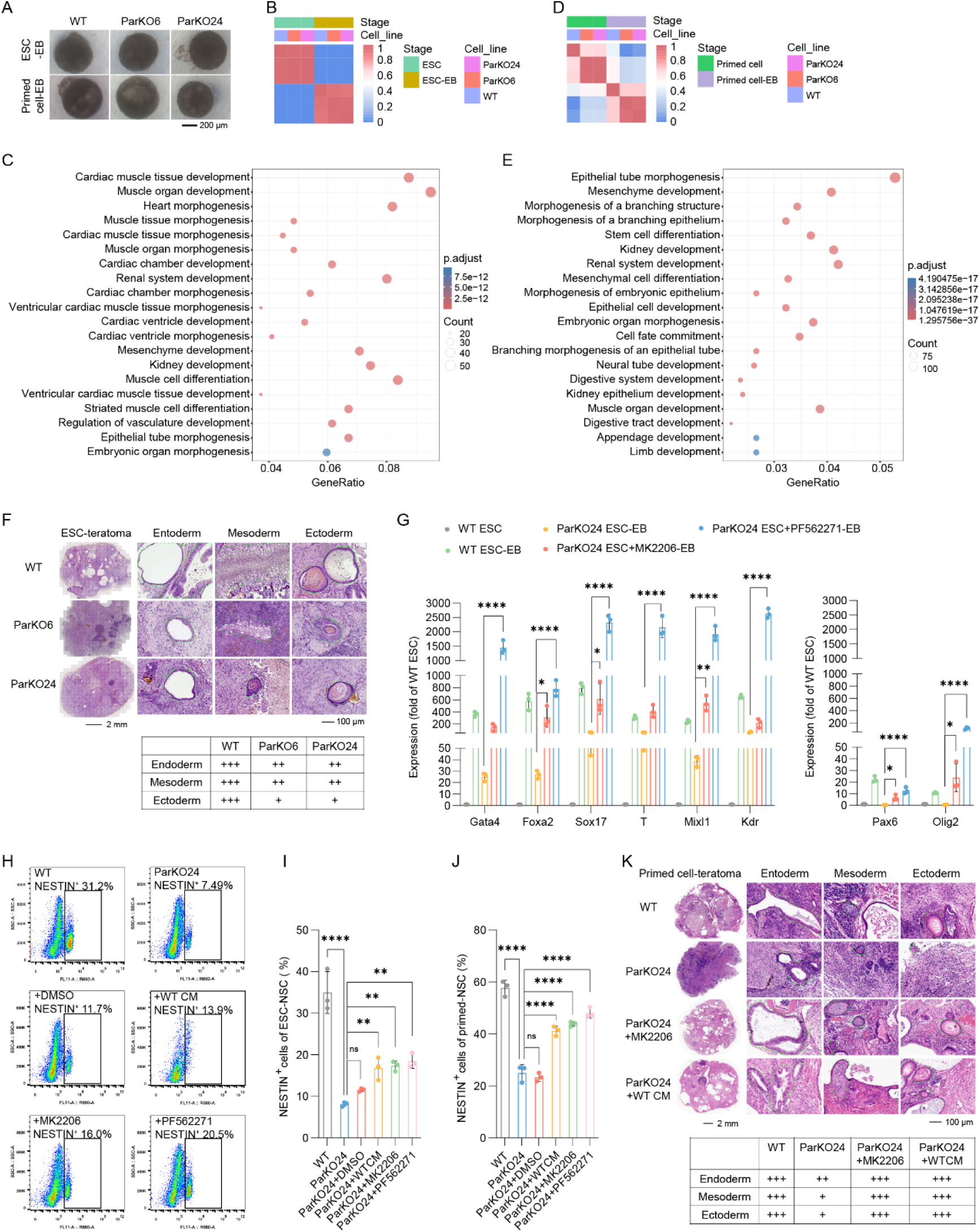
Par KO impairs lineage differentiation via the AKT–FAK signaling axis. **(A)** WT ESCs, Par KO ESCs, WT primed cells, and Par KO primed cells were subjected to EB differentiation via suspension culture. **(B-E)** Transcriptome profiling was performed on day 6 of EB differentiation separately for ESC-EBs (WT vs Par KO) and primed cell-EBs (WT vs Par KO). **(F)** Teratomas were generated by subcutaneous injection of WT and Par KO ESCs into immunodeficient mice, followed by histological assessment via H&E staining. H&E-stained sections were imaged by using the TG Tissue FAXS Plus ST. **(G)** Par KO ESCs were treated with MK2206 or PF562271 during EB differentiation, and the expression of three germ layers markers was detected. **(H-I)** Par KO ESCs were treated with MK2206 or PF562271 during the NSC differentiation. The proportion of NESTIN+ cells was detected by using the CytoFLEX. **(J)** WT, Par KO, and WT CM-, MK2206-, or PF562271-treated Par KO primed cells were differentiated into NSCs, and the proportion of NESTIN⁺ cells was quantified. **(K)** Subcutaneous injection of WT, Par KO, MK2206-or WT CM-treated Par KO primed cells into immunodeficient mice to form teratoma, followed by H&E staining for histological assessment. Experiments were repeated at least three times unless otherwise stated. Error bars represent S.D. One-way ANOVA was performed in **(G)**, (I), and **(J)**.

Transcriptomic profiling revealed significant transcriptional differences between WT and Par KO ESC-EBs (Fig. 5B). GO analysis of DEGs revealed significant enrichment in biological processes related to cardiac muscle tissue development, kidney development, epithelial tube morphogenesis, and embryonic organ morphogenesis when comparing WT with Par KO ESC-EBs (Fig. 5C). Transcriptomic divergence was already established between WT and Par KO primed cells as described previously (Fig. 2D) and persisted during EB differentiation (Fig. 5D). Consistently, DEGs identified between WT and Par KO primed cell-EBs were significantly enriched in epithelial tube morphogenesis, embryonic organ morphogenesis, cell fate commitment, and neural tube development (Fig. 5E). These results collectively demonstrate that Par KO disrupts embryonic development and lineage differentiation processes in ESCs.

To further determine the differentiation preference of Par KO ESCs, we cultured Par KO ESC-EBs for an extended period. On day 15 of EB differentiation, RT-qPCR analysis revealed significantly reduced expression of markers representative of all three germ layers in Par KO ESC-EBs compared to WT controls. These included endodermal markers (*Sox17*, *Gata4*, *Foxa2*), mesodermal markers (*Kdr*, *T*, *Mixl1*), and ectodermal markers (*Pax6*, *Olig2*) (Fig. S5A), collectively indicating impaired lineage specification capacity in the absence of Par complex.

We selected ESCs as the starting cell population for the teratoma formation assay to determine whether Par KO affects lineage differentiation *in vivo*. Both WT and Par KO ESCs formed teratomas within four weeks. However, histological analysis by H&E staining revealed that Par KO ESC-derived teratomas exhibited markedly reduced diversity and abundance of tissues representing all three germ layers (Fig. 5F). The most pronounced reduction was observed in ectodermal derivatives, particularly squamous epithelium and pigment cells. This was followed by significant decreases in endodermal lineages, including respiratory epithelium and glandular structures, as well as mesodermal derivatives such as osteoblasts.

Given the pronounced ectodermal defect, we assessed neural stem cell (NSC) differentiation. Par KO ESCs failed to form typical rosette structures (Fig. S5B), and flow cytometry showed a significant reduction in NESTIN⁺ cells compared to WT ESCs (Fig. S5C). These findings suggest that loss of the Par complex disrupts polarity-driven morphogenesis essential for lineage maturation, particularly in ectodermal derivatives. During EB differentiation, treatment with MK2206 or PF562271 robustly enhanced germ layer differentiation in Par KO ESCs (Fig. 5G and S5D). Similarly, during NSC differentiation, WT CM, MK2206, or PF562271 treatment rescued NSC morphology (Fig. S5E) and increased the proportion of NESTIN⁺ cells (Fig. 5H and 5I). Furthermore, when Par KO cells were first treated with WT CM, MK2206, or PF562271 during NPT and the resulting primed cells were then directed toward NSC differentiation, all three treatments consistently increased the proportion of NESTIN⁺ cells (Fig. 5J). For the teratoma assay, direct *in vivo* treatment with compounds or WT CM posed considerable technical challenges. We therefore adopted a pre–treatment strategy: Par KO ESCs were treated with either MK2206 or WT CM during NPT *in vitro* to generate correspondingly treated primed cells for subsequent *in vivo* teratoma formation. Both MK2206 and WT CM treatments restored tissue organization and germ layer representation in Par KO teratomas to levels comparable with WT controls, as evidenced by the robust presence of mesodermal derivatives such as osteoid tissue and ectodermal lineages including squamous epithelium (Fig. 5K).

Together, these results indicate that Par complex deficiency disrupts lineage specification and NSC generation, and that these defects can be functionally rescued by inhibiting AKT–FAK signaling.

### The AKT–FAK signaling axis mediates neural tube organoid defects in Par KO ESCs

During early embryonic development, transition from naïve ICM to primed epiblast is accompanied by the establishment of apical-basal polarity (Bedzhov & Zernicka-Goetz, 2014), which is required for apical domain changes that drive neural plate folding and neural tube closure (Matsuda et al., 2023). During neural organoid induction in vitro, naïve ESCs transition to an epiblast-like state that lies between the naïve and primed states (Park et al., 2022), indicating that neural organoid induction is closely related to NPT. Given that Par KO disrupted polarity during NPT, and impaired germ layer lineage specification of primed cells, we asked whether this defect also propagate into neural tube morphogenesis. To address this question, we derived neural tube organoids from WT and Par KO ESCs.

During the first two days of neural tube organoid induction, no significant morphological differences were observed between WT and Par KO organoids. By day 3, as ESCs transitioned toward an epiblast-like state, WT organoids developed columnar epithelial-like cells that radiated in an organized pattern from the central point. On day 4, nearly all WT organoids formed circular, sharply defined luminal structures at their centers. These lumens progressively expanded and spontaneously elongated into oval shapes. By day 6, approximately 60% of WT organoids further developed into narrow, well-defined elliptical luminal structures (Fig. 6A and 6B).

**Figure 6.**
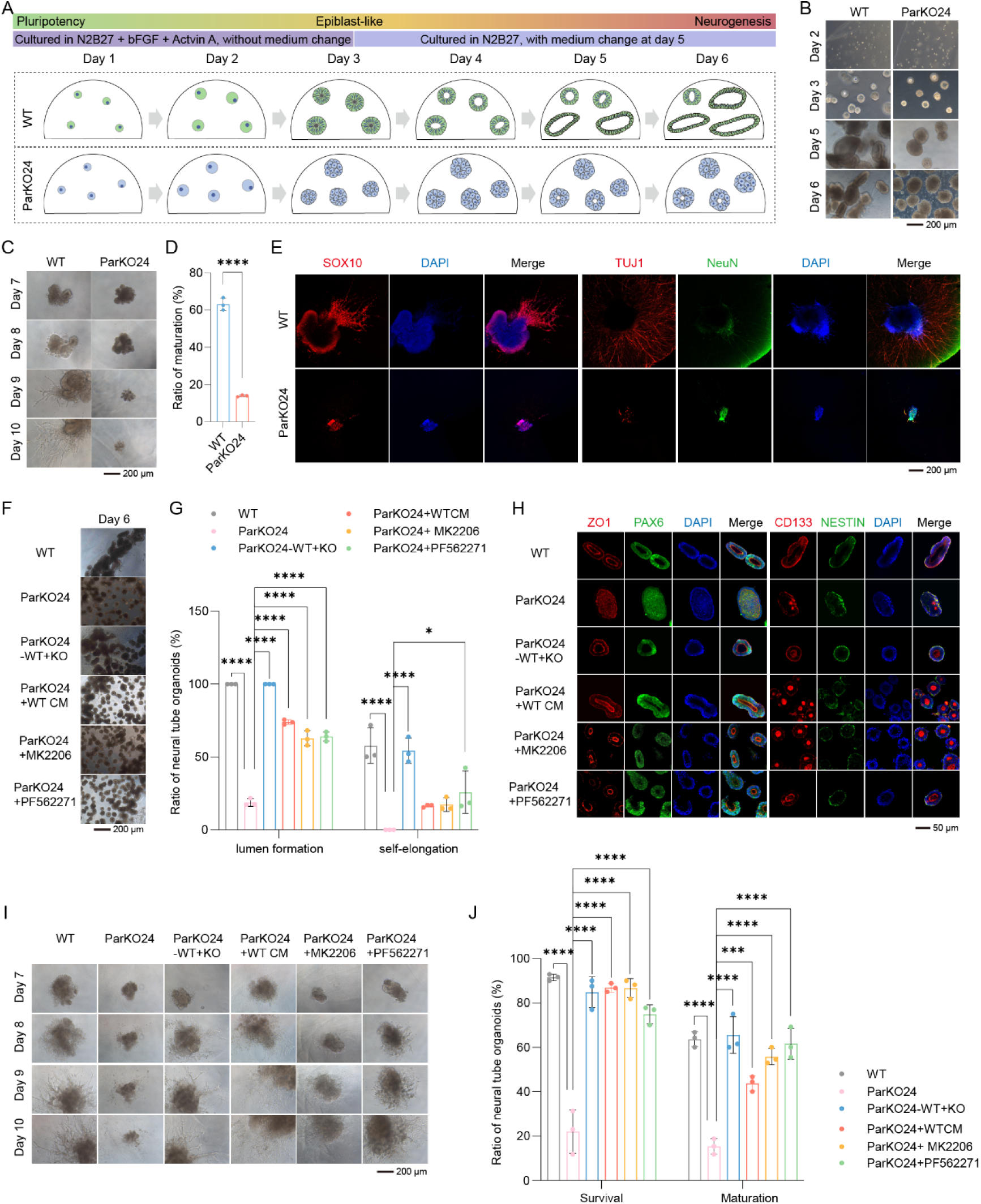
Par KO impairs neural tube organoids formation and maturation via the AKT–FAK signaling axis. **(A)** Schematic illustration of neural tube organoid induction. **(B)** The morphological changes of WT and Par KO cells during neural tube organoids induction. **(C-D)** Neural tube organoids underwent subsequent maturation culture, during which morphology was recorded **(C)** and maturation rate was analyzed **(D)**. **(E)** Mature neural tube organoids were assessed by immunofluorescence for markers of neural crest cells and neurons, such as SOX10, TUJ1, and NeuN. **(F-G)** Par KO ESCs were treated with co-culture, WT CM, MK2206, or PF562271 during neural tube organoid induction, the lumen formation and spontaneous elongation efficiency under different conditions were assessed. **(H)** After treatment with co-culture, WT CM, MK2206, and PF562271, the expression of the lumen markers (ZO1 and CD133) as well as the NSC markers (PAX6 and NESTIN) was detected in Par KO neural tube organoids. **(I-J)** Par KO ESCs were treated with co-culture, WT CM, MK2206, or PF562271 during neural tube organoid maturation culture, the survival and maturation efficiency under different conditions were assessed. Experiments were repeated at least three times unless otherwise stated. Error bars represent S.D. One-way ANOVA was performed in **(D)**, **(G)**, and **(J)**.

In contrast, Par KO organoids failed to form columnar cellular structures and displayed disorganized cell alignment without radial symmetry on day 3. By day 4, these organoids exhibited only volume increase without forming typical lumens. By day 5, only a small fraction of Par KO organoids (∼18%) formed lumen-like structures; however, these structures were small, circular, poorly delineated, and failed to elongate. By day 6, these aberrant lumens showed no further expansion or elongation (Fig. 6A and 6B).

During subsequent maturation culture (days 7–10), approximately 63% of WT neural tube organoids differentiated into numerous cells with neuronal axons. In contrast, Par KO organoids exhibited substantial apoptosis, with only about 13% developing axon-bearing cells (Fig. 6C and 6D). Immunofluorescence analysis for neural crest cell markers (SOX10), early neuron markers (TUJ1), and mature neuron markers (NeuN) revealed abundant neural crest cells and neurons in WT neural tube organoids, whereas Par KO organoids showed significantly reduced numbers of these cell types (Fig. 6E). These results demonstrate that loss of the Par complex severely disrupts the differentiation and maturation of ESCs into neural tube.

We next examined whether the morphogenetic defects in Par KO neural tube organoids could be rescued by co-culture with WT ESCs, WT CM, MK2206 or PF562271. All four treatments significantly enhanced both lumen formation efficiency and the frequency of spontaneous elongation by day 6 (Fig. 6F and 6G). Immunofluorescence analysis was performed to assess the localization of apical markers (ZO1, CD133) and the expression of NSC markers (PAX6, NESTIN). In untreated Par KO organoids, while PAX6 and NESTIN were expressed, ZO1 and CD133 showed aberrant localization with loss of apical polarity. All rescue treatments restored the apical distribution of ZO1 and CD133 to a pattern resembling that of WT organoids (Fig. 6H). During maturation culture, the four treatments, including co-culture with WT ESCs, WT CM, MK2206 or PF562271, also promoted the survival and maturation efficiency of Par KO organoids (Fig. 6I and 6J).

Collectively, these findings identify a critical role for the Par complex in maintaining apical-basal polarity and promoting lumenogenesis during neural tube morphogenesis. The successful rescue of these morphological defects, which was achieved through co-culture with WT cells, WT CM, MK2206, or PF562271, confirms the functional centrality of the FAK signaling pathway downstream of Par complex activity.

## Discussion

Cell polarity establishment marks a critical transition for embryos to initiate lineage specification, a process governed by conserved polarity complexes (Etemad-Moghadam et al., 1995; Wang et al., 2018). Here, by generating Crumbs, Par, or Scrib complex knockout cells, we identified a specific and essential role for the Par complex during NPT. Par deficient cells failed to adopt the characteristic flat monolayer morphology of WT primed cells, highlighting its dominant function in morphological organization. This aligns with studies showing that targeted Par complex localization can induce polarity de novo (Watson et al., 2023). Further analysis revealed that Par loss disrupted apical junctional integrity, as evidenced by mislocalization of the adherens junction marker CDH1 and the tight junction marker ZO1. This supports the established role of the Par complex in forming and stabilizing both junction types (Hirose et al., 2002). Functionally, Par KO impaired differentiation across all three germ layers in EB and teratoma assays, and specifically disrupted NSC and neural tube organoid formation. The profound differentiation defect likely stems from the fundamental role of the Par complex in establishing polarity of various cell types during embryonic development, as exemplified by the finding that even a single amino acid mutation in *Par3* is sufficient to cause neural tube defects (Chen et al., 2017). Collectively, our results establish the Par complex as a master regulator of ESC differentiation potential, providing a mechanistic basis for the early embryonic lethality observed upon its deletion (Alarcon, 2010).

The establishment of epithelial cell polarity critically depends on the ECM, a network primarily composed of collagen, laminin, and glycoproteins that regulates cell adhesion and provides spatial cues through integrin-mediated signaling (Sun et al., 2019). Our transcriptomic analysis revealed that Par KO disrupts this axis, with dysregulation of genes involved in ECM-receptor interaction, focal adhesion, and cytoskeleton organization during NPT. Both AKT and FAK inhibition rescued the flat monolayer morphology in Par KO primed cells. Notably, AKT inhibition reduced FAK phosphorylation, positioning FAK as a key downstream effector in maintaining cell morphology. This is consistent with the known ability of FAK to regulate cell adhesion and ECM-cell interaction (Dawson et al., 2021). This regulatory node was further highlighted by the rescue achieved with WT CM, which also normalized p-FAK levels, indicating that ECM, integrin, and FAK signaling together regulate cell behavior (Ilic et al., 1995). Proteomic analysis identified differential expression of proteins involved in cell adhesion and integrin signaling between WT and Par KO CM. Transcriptional profiling further revealed that genes commonly upregulated in MK2206-treated Par KO cells, WT CM-treated Par KO cells, and WT cells were enriched in pathways governing cell adhesion and ECM organization. Among the candidate factors, LEFTY1/2 stood out for their strong rescue capability and their higher abundance in WT CM versus Par KO CM, suggesting LEFTY acts as a critical extracellular mediator supplied by WT cells. LEFTY, a morphogen for left–right (L-R) determination (Duboc et al., 2008; Meno et al., 1996), regulates embryonic processes such as gastrulation (Branford & Yost, 2002; Dai et al., 2016; Feldman et al., 2002), and plays a key role in tissue and organoid formation *in vitro* (Song et al., 2021). Moreover, previous studies have shown that LEFTY regulates Par complex function through TGFβ/BMP signaling during neural tube morphogenesis (Amarnath & Agarwala, 2016), whereas our findings demonstrate that Par complex regulates LEFTY via AKT signaling. Therefore, the interplay between Par complex and LEFTY in the regulation of embryonic development warrants further investigation.

Notably, although Par KO primed cells are dome-shaped, the expression of primed markers was comparable to WT cells. This implies that Par KO leads to pronounced defects in cell morphology, rather than blocking cells differentiation from naïve to primed state, resulting in cells that reach a primed state with aberrant dome-shaped morphology. As transitions in cell potency are frequently associated with coordinated morphogenetic changes (Endoh & Niwa, 2022), the morphological changes in Par KO primed cells may be linked to their altered differentiation potential. In line with this, the functional importance of Par complex extends beyond morphology to differentiation potential: EB differentiation, teratoma formation, and directed NSC differentiation. Inhibition of FAK signaling restored the impaired differentiation potential of Par KO cells across these assays This demonstrates that Par complex governs differentiation capacity through a conserved, morphology-linked mechanism. WT CM was omitted from EB differentiation assays due to volume constraints in the suspension culture system. Notably, we observed a nuanced interaction in co-culture system: while WT cells rescued the morphological defects, they simultaneously suppressed proliferation of Par KO cells (Fig. 3D and 3E). This suggests the activation of cell competition, a quality-control process that may selectively eliminate suboptimal cells during development (Di Gregorio et al., 2016; Hashimoto & Sasaki, 2020; Maruyama & Fujita, 2022). Collectively, our findings delineate a signaling pathway in which the Par complex governs cell polarity establishment by sequentially modulating AKT activity, FURIN-mediated LEFTY maturation, and ultimately FAK phosphorylation. However, our findings are limited to *in vitro* systems, and it remains unclear whether the Par–AKT–FURIN–LEFTY–FAK axis operates similarly *in vivo*. Par disruption impairs neuroepithelial polarity and tissue morphogenesis (Zhang & Wei, 2022), while AKT, FURIN, LEFTY, and FAK participate in developmental signaling (Ding et al., 2021; Liang et al., 2022; Mamali et al., 2007; Nozaki et al., 2025; Zhang et al., 2008). Consistent with this, FURIN deficiency causes defects in embryonic turning and heart looping and alters LEFTY expression (Constam & Robertson, 2000), and LEFTY in turn modulates FAK phosphorylation (Alowayed et al., 2016). Nevertheless, whether Par coordinates these pathways remains unclear.

In vertebrate embryos, neuroepithelial cells divide predominantly along the apical-basal axis, this polarity is essential for neural tube formation (Herrera et al., 2021). During neural plate folding, epithelial polarization enables neural plate bending, neural fold convergence, and tube closure (Shi, 2022), with polarity proteins mediating intercellular adhesion to maintain structural integrity (Hong et al., 2010; Legere et al., 2024). Neural tube defects (NTDs), including anencephaly caused by *Pard3* deletion, represent severe developmental disorders with high mortality and disability rates, often stemming from failure of embryonic neural tube closure (Chen et al., 2017; Nychyk et al., 2022). Although the molecular basis of NTDs remains incompletely defined, polarity defects in neuroepithelial cells represent a major contributing factor (MacGowan et al., 2025). The Par complex plays a key regulatory role in apical-basal axis establishment during neural tube formation (Zhang & Wei, 2022; Zovein et al., 2010). In mice, *Pard3* deficiency causes neural developmental abnormalities and mid-gestational embryonic lethality (Hirose et al., 2006). Recent studies have shown that Par complex abnormalities disrupt apical tight junctions and neuroepithelial polarization, contributing to NTDs in humans (Chen et al., 2017; Gao et al., 2012). Consistent with these findings, using ESC-derived neural tube organoids, we demonstrated that Par complex deficiency significantly reduces lumen structure formation efficiency and impairs spontaneous elongation. During maturation, Par KO neural tube organoids exhibited increased apoptosis and reduced maturation efficiency, underscoring the critical role of the Par complex in neural tube formation. Given that we observed morphological defects in ParKO primed cells, it is reasonable to speculate that neural tube defects caused by Par KO may occur as early as the primed state. Treatment with WT CM, MK2206, or PF562271 significantly improved lumen formation and maturation efficiency, with PF562271 showing the most prominent rescue effect. These results suggest FAK phosphorylation as a downstream mediator of Par complex-regulated ESC differentiation and neural tube formation. This insight not only deepens our understanding of developmental plasticity but also suggests promising therapeutic strategies for NTDs.

## Materials and Methods

### Animal model

Female BALB/C Nude mice were purchased from GemPharmatech and maintained in ventilated cages under a 12:12 light:dark cycle at a controlled temperature, having *ad libitum* access to standard food and water. Five-week-old mice were used in this paper for the subcutaneous tumor formation assay, and maintained in the same conditions for 4 weeks. The experimental protocol complies with the requirements of animal welfare ethics, has been approved by the animal experimental ethics review in Guangzhou Medical University, and is permitted to be carried out, with the ethics review number N2025-25023.

### Cell culture

OG2 embryonic stem cells (egfp reporter genes driven by the Oct4 promoter, GOF18ΔPE, CBA/CaJ X C57BL/6J) (Szabó et al., 2002) used in this study were recovered from liquid nitrogen, and cultured in mES medium (containing DMEM/F12, Neurobasal, 1% GlutaMAX, 1% NEAA, 1% Sodium Pyruvate, 0.5% N2 supplement, 1% B27 supplement, 0.1 mM β-Mercaptoethanol, 1000 units/mL LIF, 3 μM CHIR99021, and 1 μM PD0325901). During passage, ESCs were digested with Accutase at room temperature for 2 minutes, then terminated by DMEM/F12.

Primed cells were derived from NPT induction, and cultured in NPT medium (containing DMEM/F12, Neurobasal, 1% GlutaMAX, 1% NEAA, 0.5% N2 supplement, 1% B27 supplement, 0.1 mM β-Mercaptoethanol, 20 ng/mL bFGF, 20 ng/mL Activin A, 2 μM XAV939, and 3 μM CHIR99021). During passage, primed cells were digested with Accutase at room temperature for 3 minutes, then terminated by DMEM/F12. After centrifugation at 250 × *g* for 5 minutes, the cell clusters were resuspended in fresh NPT medium and gently shaken to break them into small clumps without dispersing them into single cells. An appropriate number of cell clusters was transferred to a petri dish and placed in an incubator.

293T cells were purchased from ATCC (www.atcc.org), and cultured in MEF medium (containing DMEM/High Glucose, 10% FBS, 1% GlutaMAX, and 1% NEAA). During passage, 293T cells were digested with 0.25% Trypsin-EDTA at 37 °C for 1 minute, then terminated by MEF medium.

All cell lines were cultured at 37 °C in a humidified incubator with 5% CO₂, and routinely tested to exclude mycoplasma contamination.

### Construct knockout cell lines

Specific sgRNA sequences targeting the corresponding genes were designed and cloned into the lentiCRISPRv2 vector to construct recombinant knockout plasmids (Joung et al., 2017). The recombinant lentiviruses were produced using a PEI-mediated lentiviral packaging system, and the viral supernatants were collected to infect ESCs. Following two rounds of viral infection, the ESCs were subjected to Puromycin selection for 1 day to obtain stably infected cell populations. The surviving ESCs were digested into single-cell suspensions, and monoclonal cell lines were isolated using the 96-well plate limiting dilution method. After expansion of the monoclonal ESCs, genotype sequencing and western bloting were performed to identify and select positive clones with complete knockout of the target gene for subsequent experiments. All sgRNA sequences were listed in the Key Resources Table.

### Naïve-to-primed transition (NPT)

ESCs were digested into single cells with Accutase and seeded into gelatin-precoated culture dishes at a density of 10,000 cells/cm². They were cultured in mES medium until cell adhesion (approximately 12 hours), then mES medium was replaced with NPT medium for continuous induction over 6 days with fresh medium replaced daily. During the NPT process, the percentage of round morphology was quantified based on both the number and area of cell clusters. In the first two days of NPT, individual Par KO ESCs progressively developed into separated, flat cell clusters. By day 3, these flat clusters began to contract, aggregate, and form domed colonies. Thus, the percentage of round morphology was calculated as the number of domed colonies divided by the total number of cell clusters. However, from day 3 to day 6, when adjacent flat clusters merged into continuous sheets, the number of flat clusters was estimated based on area. Specifically, using half the average distance between adjacent domed colonies as the radius, a circular area was defined to represent the typical area of a single flat cluster. This circular reference was then used to partition the merged flat regions and estimate the count of individual flat clusters.

### ESC-to-neural stem cell (NSC) differentiation

ESCs were digested into single cells with Accutase and seeded into Matrigel-precoated culture dishes at a density of 10,000 cells/cm².They were cultured in mES medium until cell adhesion (approximately 8 hours), then mES medium was replaced with NSC medium (containing DMEM/F12, 1% N2 supplement, 2% B27 supplement). NSC differentiation was continued for 7 consecutive days, with fresh NSC medium replaced daily during this period. Matrigel (CORNING, 356234) was diluted with DMEM/F12 at a 1:1 ratio.

### ESC-to-embryoid body (EB) differentiation

ESCs were digested into single cells with Accutase and resuspended in mES-lif-2i medium (containing DMEM/High Glucose, 15% FBS (Gibco, 1099141C), 1% GlutaMAX, 1% NEAA, 1% Sodium Pyruvate, 0.1 mM β-Mercaptoethanol), then diluted to a density of 20,000 cells/mL. After mixing thoroughly, 2 mL of the cell suspension was vertically and evenly dropped onto the inner side of the 10 cm² culture dish lid using multichannel pipette with 20 μL/drop. Subsequently, 3 mL PBS was added to the 10 cm² culture dish, and the lid with cell droplets was gently and quickly placed back on the dish.The dish was transferred to an incubator for hanging drop culture for 3 days, with no medium change during this period. On Day 3, EBs were collected into a centrifuge tube using a Pasteur pipette and allowed to settle naturally for approximately 5 minutes until the EBs precipitated to the bottom of the tube. The EBs were then gently transferred to a T25 flask containing 8 mL fresh mES-lif-2i medium. The flask was then transferred to a horizontal shaker inside the humidified incubator and cultured continuously for another 6 days at 60 RPM, 37 °C, and 5% CO₂, with 1 mL of fresh mES-lif-2i medium supplemented daily.

### Teratoma formation assay in immunodeficient mice

A total of 1×10⁶ ESCs were transferred to a 1.5 mL EP tube with 1 mL DPBS, centrifuged to remove the DPBS, and the tube was kept on ice for subsequent operations. The ESCs were immediately resuspended in 100 μL of a 1:1 mixture of DMEM/F12 and Matrigel (CORNING, CLS356234), then subcutaneously injected into the dorsal flank of immunodeficient mice via an insulin syringe. The mice were maintained in ventilated cages under a 12:12 light:dark cycle at a controlled temperature, having *ad libitum* access to standard food and water. After 4 weeks, the teratoma tissues were harvested, then processed for sectioning and hematoxylin-eosin (HE) staining. HE-stained sections were imaged by using the TG Tissue FAXS Plus ST (TissueGnostics, Austria), and the scanned images were analysed using TissueFAXS Viewer software (version 7.1.6245.142).

### ESC-to-neural tube organoid differentiation

The method of neural tube organoid induction was performed as previously reported (Park et al., 2022). The ESCs were digested into single cells with Accutase and resuspended in N2B27 medium (containing DMEM/F12, Neurobasal, 0.5% GlutaMAX, 1% NEAA, 1% Sodium Pyruvate, 0.5% N2 supplement, 1% B27 supplement, 0.1 mM β-Mercaptoethanol). A total of 1200 cells were transferred to a new EP tube containing 300 μL of N2B27 and centrifuged at 250 × *g* for 5 minutes, then embedded in 10 μL of Matrigel (CORNING, CLS354248) at 1200 cells per drop. The 10 μL cell suspension was dropped into one well of a 24-well plate (Corning) containing 500 μL of Epi medium (N2B27 supplemented with 12 ng/mL bFGF and 20 ng/mL Activin A), and cultured for 3 days without medium change. On day 3, the Epi medium was replaced with 1 mL of N2B27 medium, and the organoids were cultured for another 2 days. On day 5, the medium was replaced with 1 mL of fresh N2B27 medium, and the organoids were cultured until day 6. For maturation of neural tube organoid, the neural tube organoids in Matrigel drop on day 6 were gently dissolved with 500 μL mixture of 3 mg/mL Collagenase IV and 25 U dispase (both dissolved in HBSS), then the released organoids were re-embedded in Matrigel at a concentration of 1 organoid/10 μL and dropped into each well of a 96-well plate with glass bottom. 100 μL of N2B27 medium was added to each well. The organoids were further cultured until day 10, with the medium changed daily.

### Immunostaining and confocal microscopy

For cellular immunofluorescence staining, cells were fixed in 4% paraformaldehyde (PFA) at room temperature for 30 minutes, then blocked in QuickBlock™ Blocking Buffer for Immunol Staining (Beyotime Biotechnology, P0260) at 37 °C for 30 minutes. Subsequently, primary antibodies and secondary antibodies were incubated sequentially. Primary antibodies were diluted at 1:200 in QuickBlock™ Primary Antibody Dilution Buffer for Immunol Staining (Beyotime Biotechnology, P0262), and incubated on a shaker at 4°C overnight. Secondary antibodies were diluted at 1:500 in QuickBlock™ Secondary Antibody Dilution Buffer for Immunofluorescence (Beyotime Biotechnology, P0265), and incubated on a shaker at room temperature for 1 hour (protect from light hereafter). The nuclei were counterstained with 4’,6-diamidino-2-phenylindole (DAPI). Confocal images of cells were captured using the Zeiss LSM 800, and the scanned images were analysed using ZEN software (ZEN 2012).

### Flow cytometry analysis

Cells were harvested and fixed in 4% PFA, then permeabilized and blocked at room temperature for 30 minutes. Permeabilization and blocking solution contained PBS, 0.1% Triton X-100 and 1% FBS. Incubated primary antibody at 4 °C for 2 hours and secondary antibody at room temperature in the dark for 45 minutes. Primary antibody and secondary antibody were diluted to 1:200 and 1:500, respectively. After staining, cells were resuspended at an appropriate volume of PBS according to the cell number. The cells were filtered through a 300-mesh filter and analyzed using the CytoFLEX flow cytometer (BECKMAN COULTER).

### RNA-seq and RT-quantitative PCR (RT-qPCR)

Samples were collected as this study described and lysed in VeZol Reagent (Vazyme, R411-02) before being sent to sequencing company (Guangzhou HeQin Biotechnology Co., Ltd.) for RNA-seq. For qPCR, total RNA was extracted from samples using VeZol Reagent, and 1 μg of RNA was reverse-transcribed into cDNA using HiScript IV All-in One Ultra RT SuperMix for qPCR (Vazyme, R433-01). Then the transcript levels of the genes were quantified using ChamQ Blue Universal SYBR qPCR Master Mix (Vazyme, Q312-03) on a CFX-96 Real-Time system (CFX96, Bio-Rad). Primers used in this study are listed in Key Resource Table.

### Immunoblotting

Proteins were separated by SDS-PAGE gel and transferred to PVDF membranes. Following this, the membranes were blocked for 1 hour at room temperature using 5% non-fat milk in TBST. The membranes were then probed with the primary antibodies overnight at 4°C. Incubated the secondary antibodies for 1 hour at room temperature, then imaged with Immobilon Western Chemiluminescent HRP Substrate on Chemiluminescent/ Fluorescent/ Gel Imaging and Analysis System (ChampChemi610Plus, SINSAGE).

### Quantification and statistical analysis

All statistical analyses were performed using GraphPad Prism 8. Independent replicates: n ≥ 3.

Data are presented as means ± SD unless otherwise stated. Statistical comparisons among multiple groups were determined by One-way ANOVA, correct for multiple comparisons using statistical hypothesis testing with Tukey. while comparisons between two individual groups were conducted using Student’s t-test, unless otherwise stated. The statistical analysis criteria were: * P < 0.05, ** P < 0.01, *** P < 0.001, **** P < 0.0001.

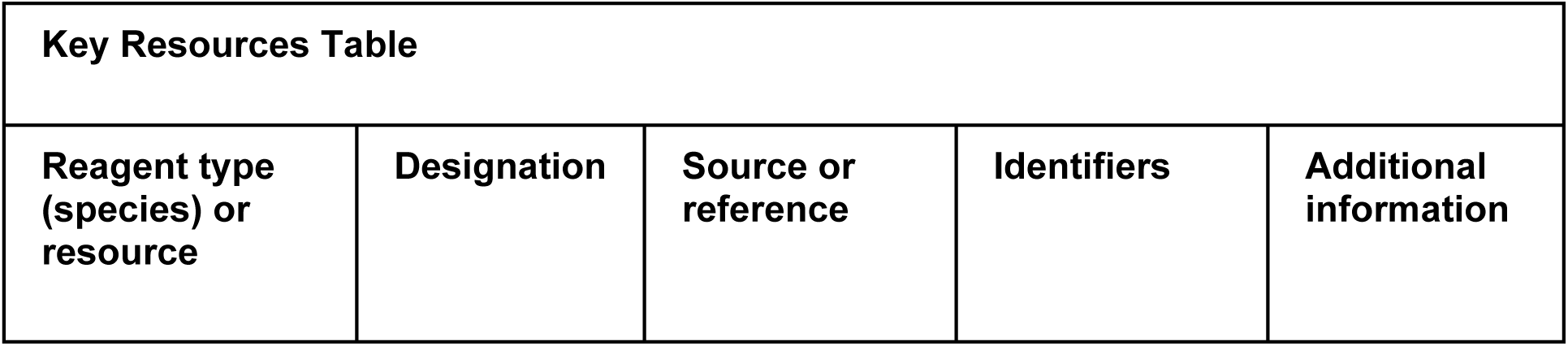
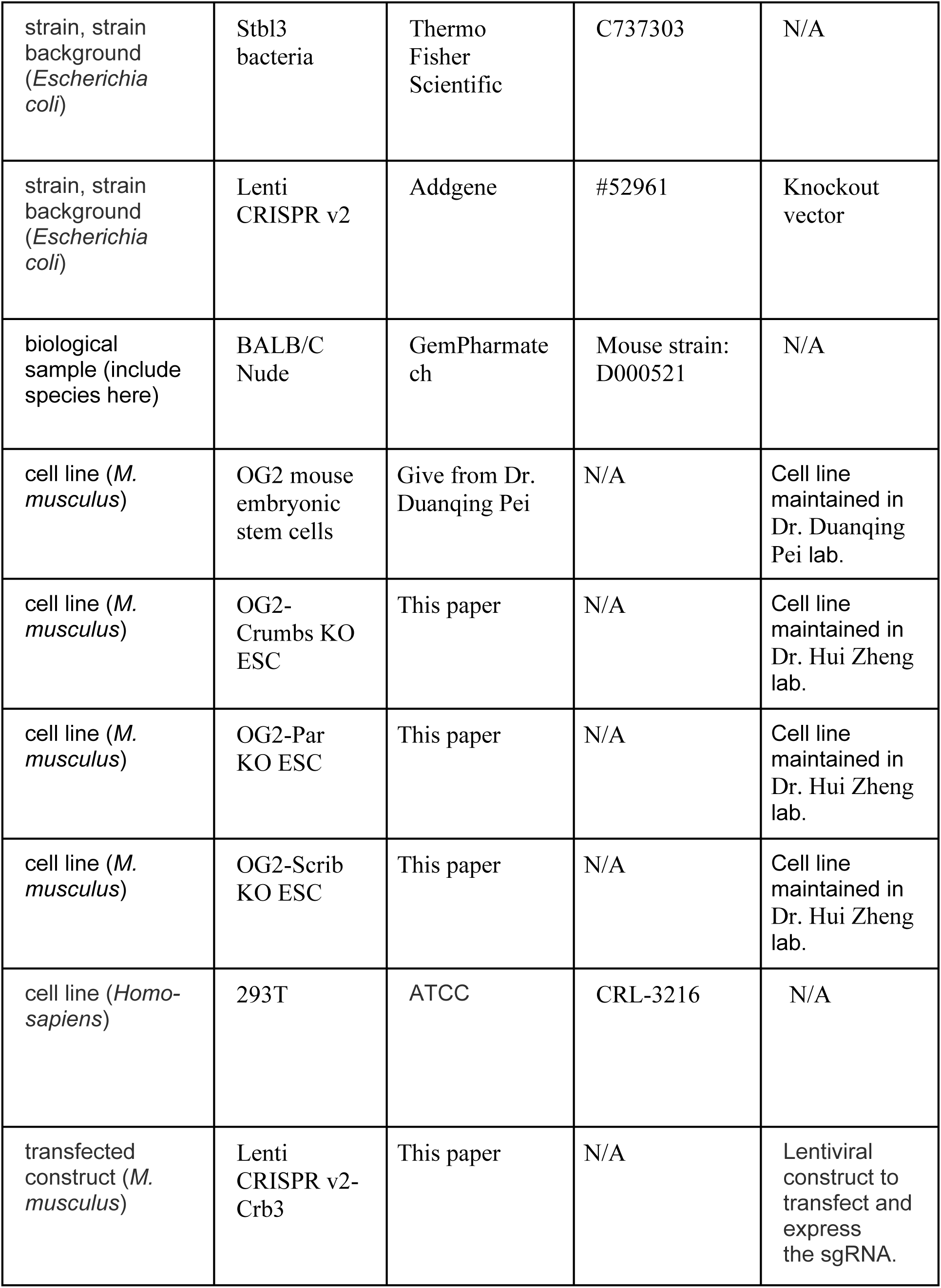
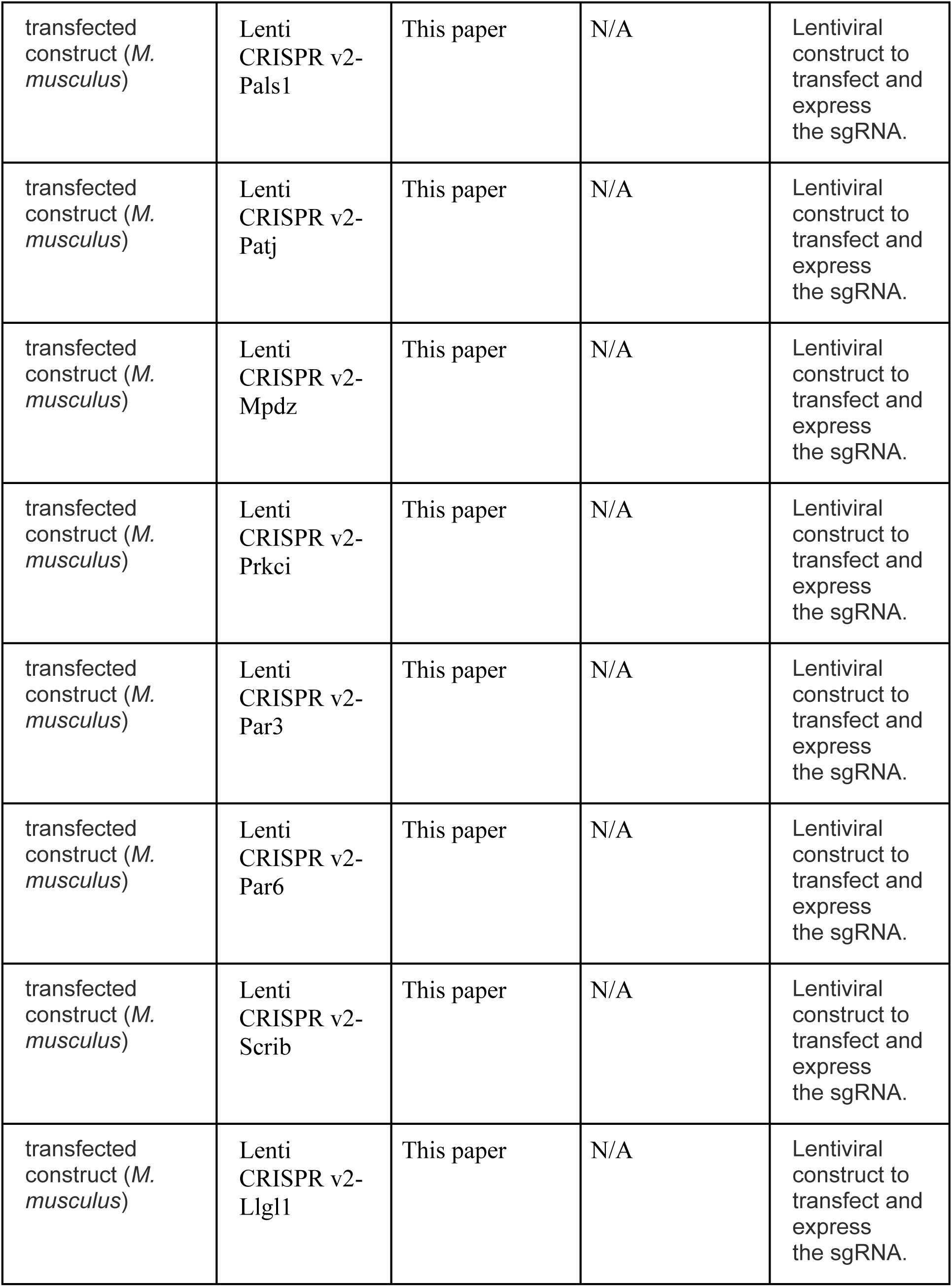
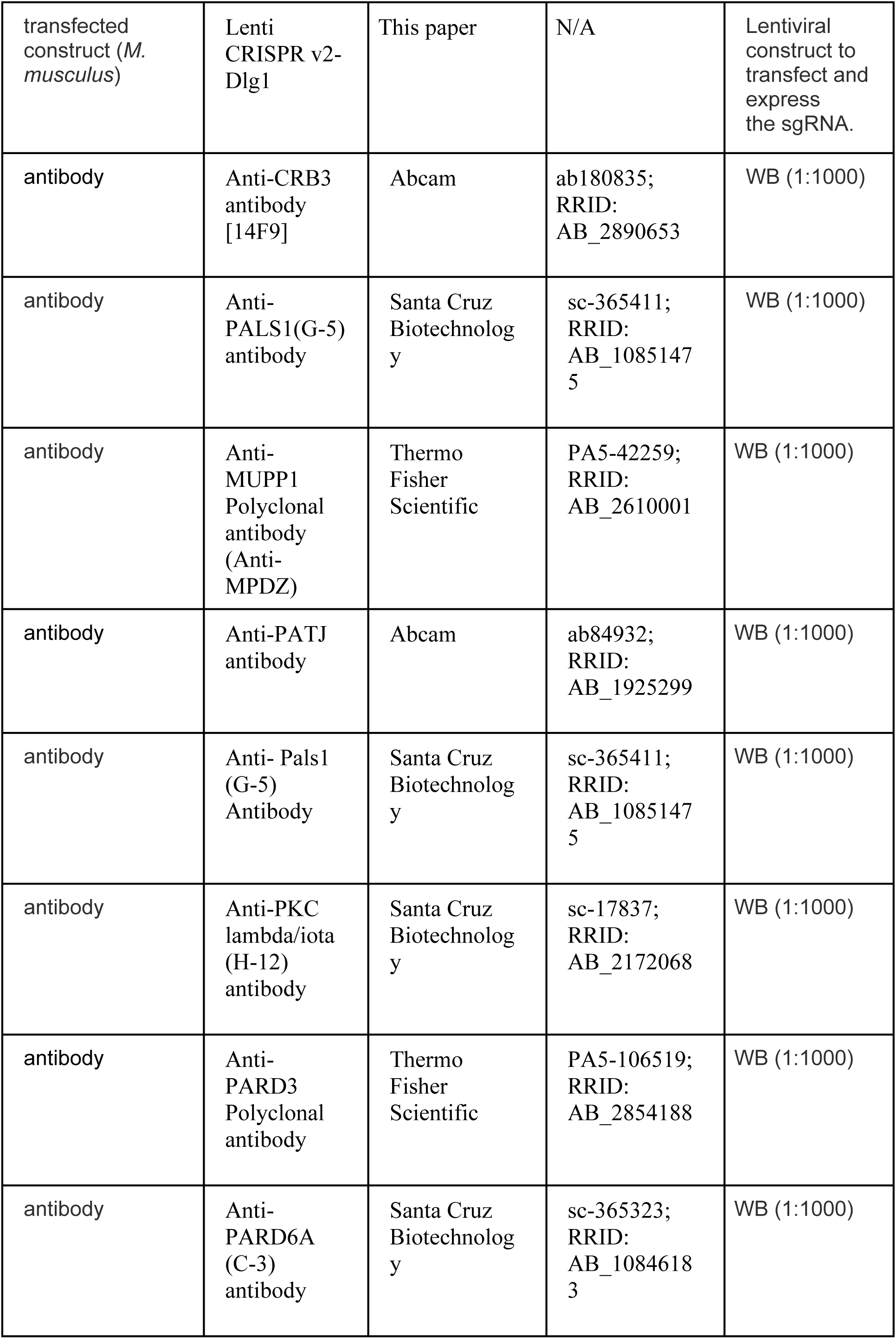
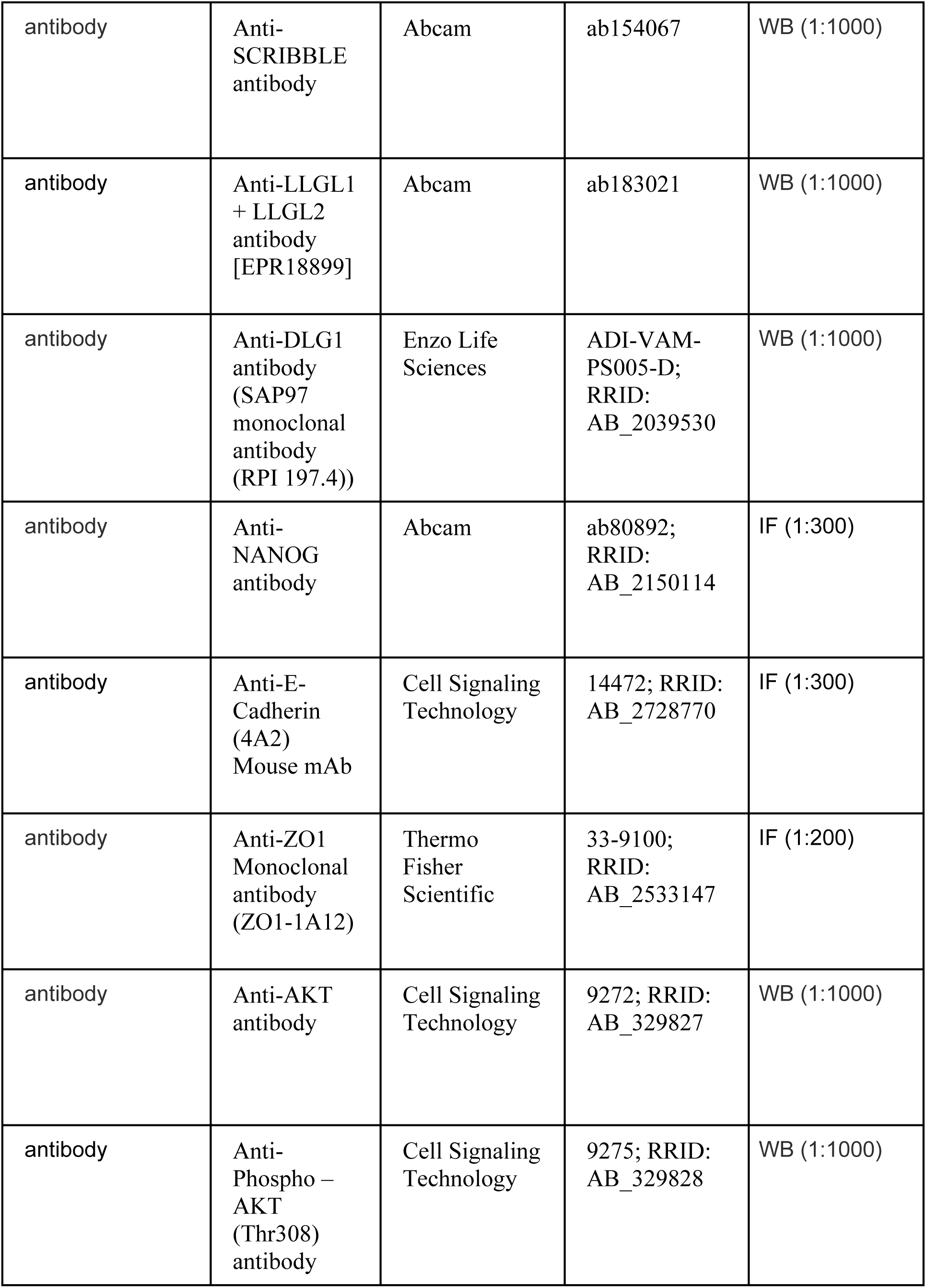
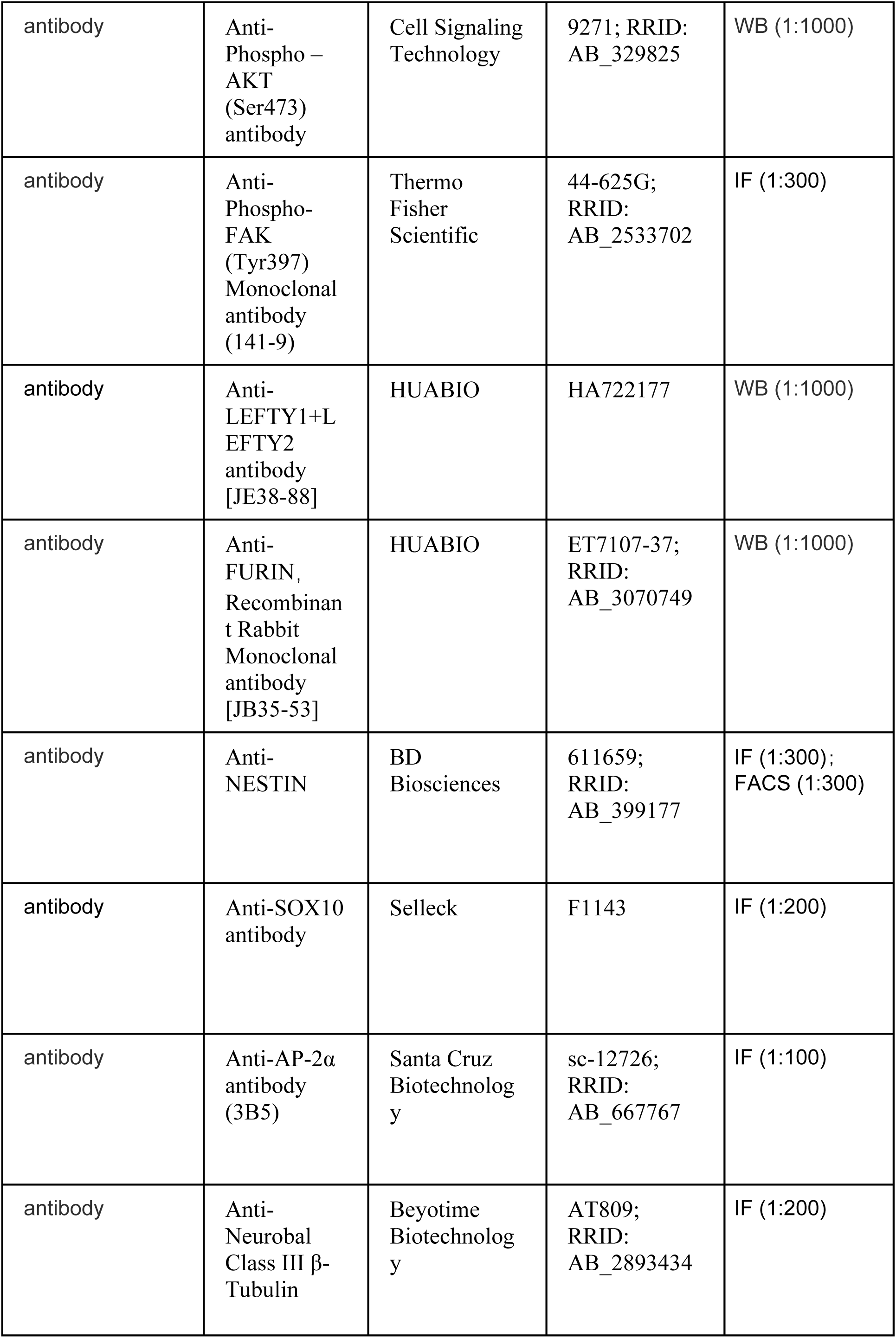
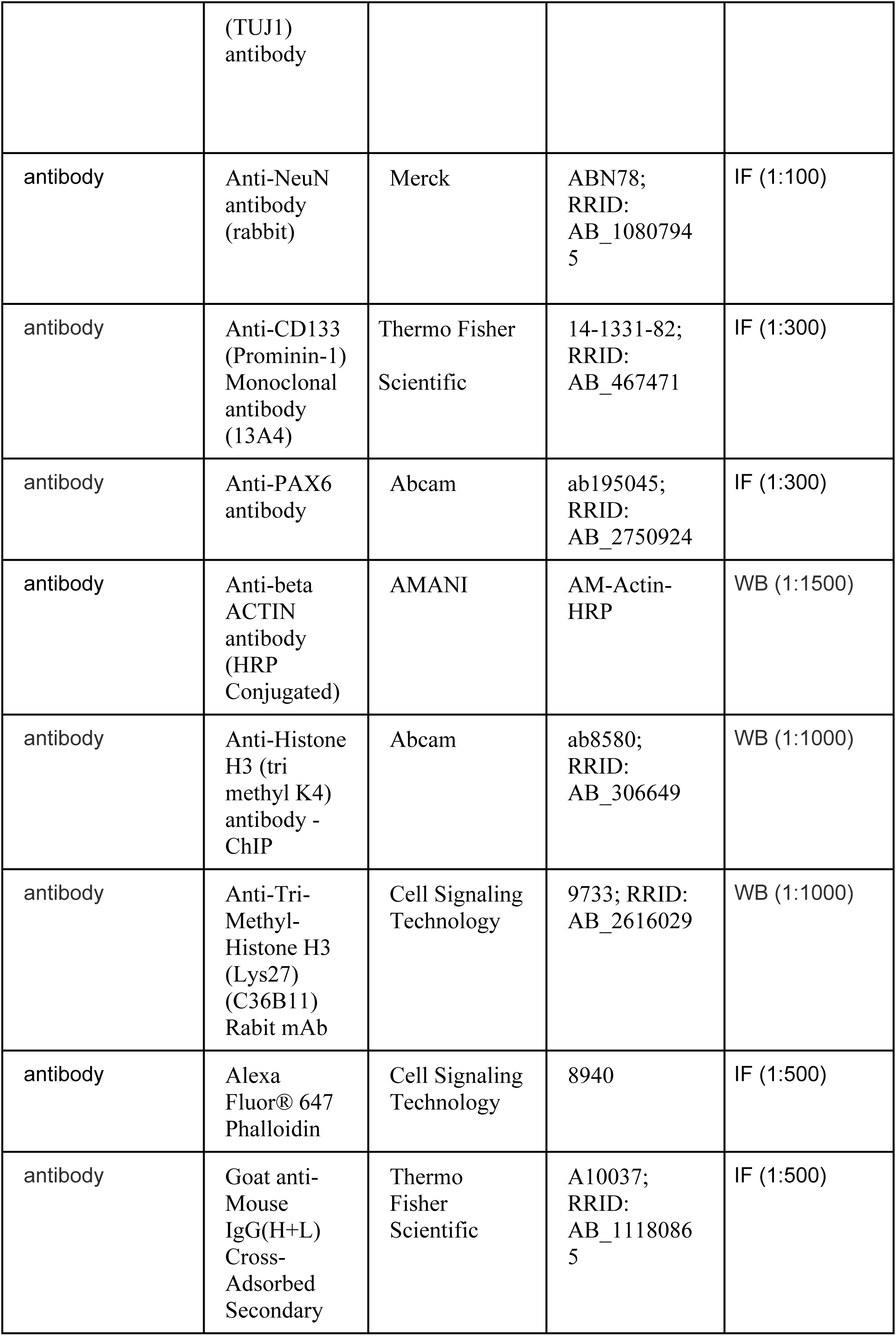
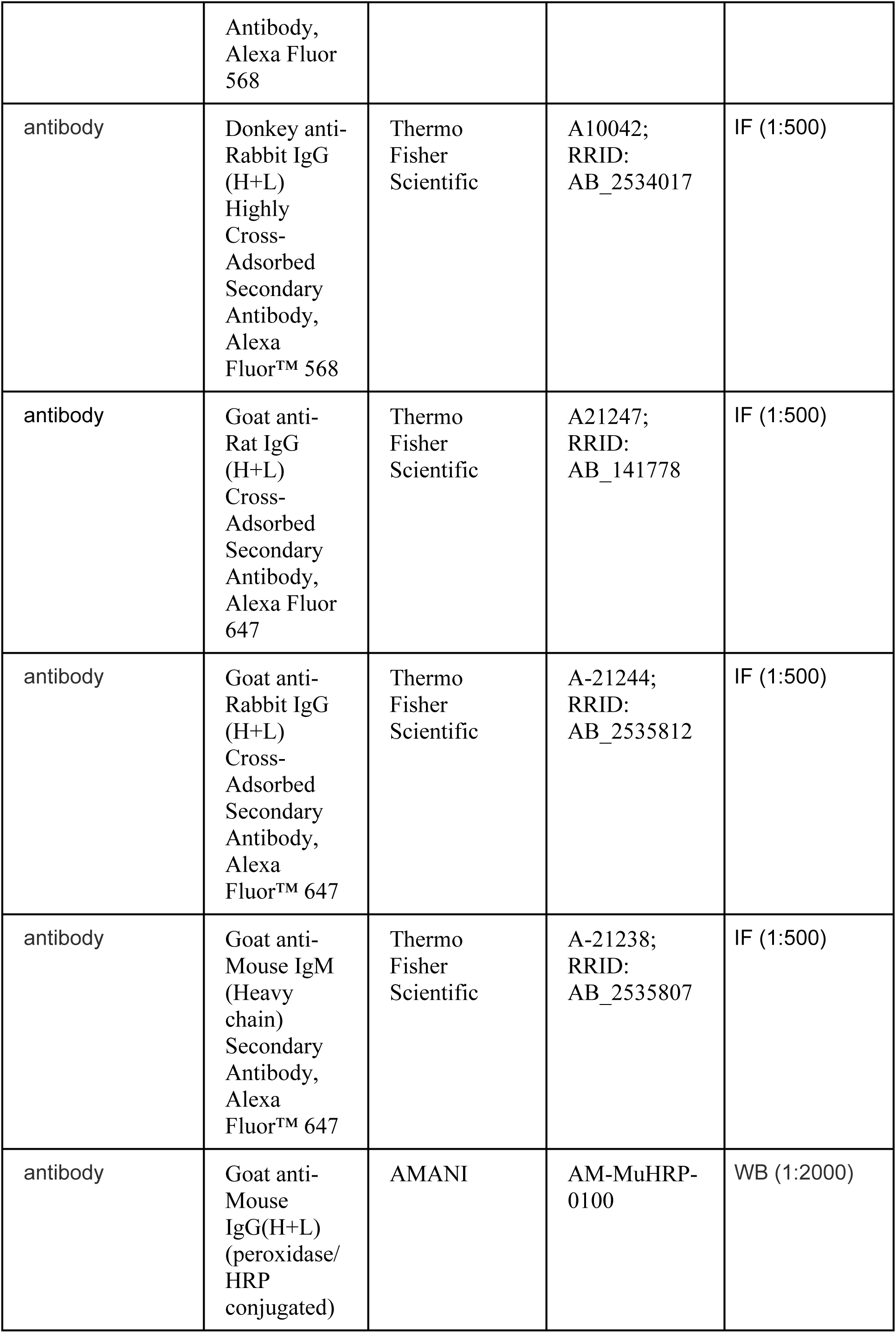
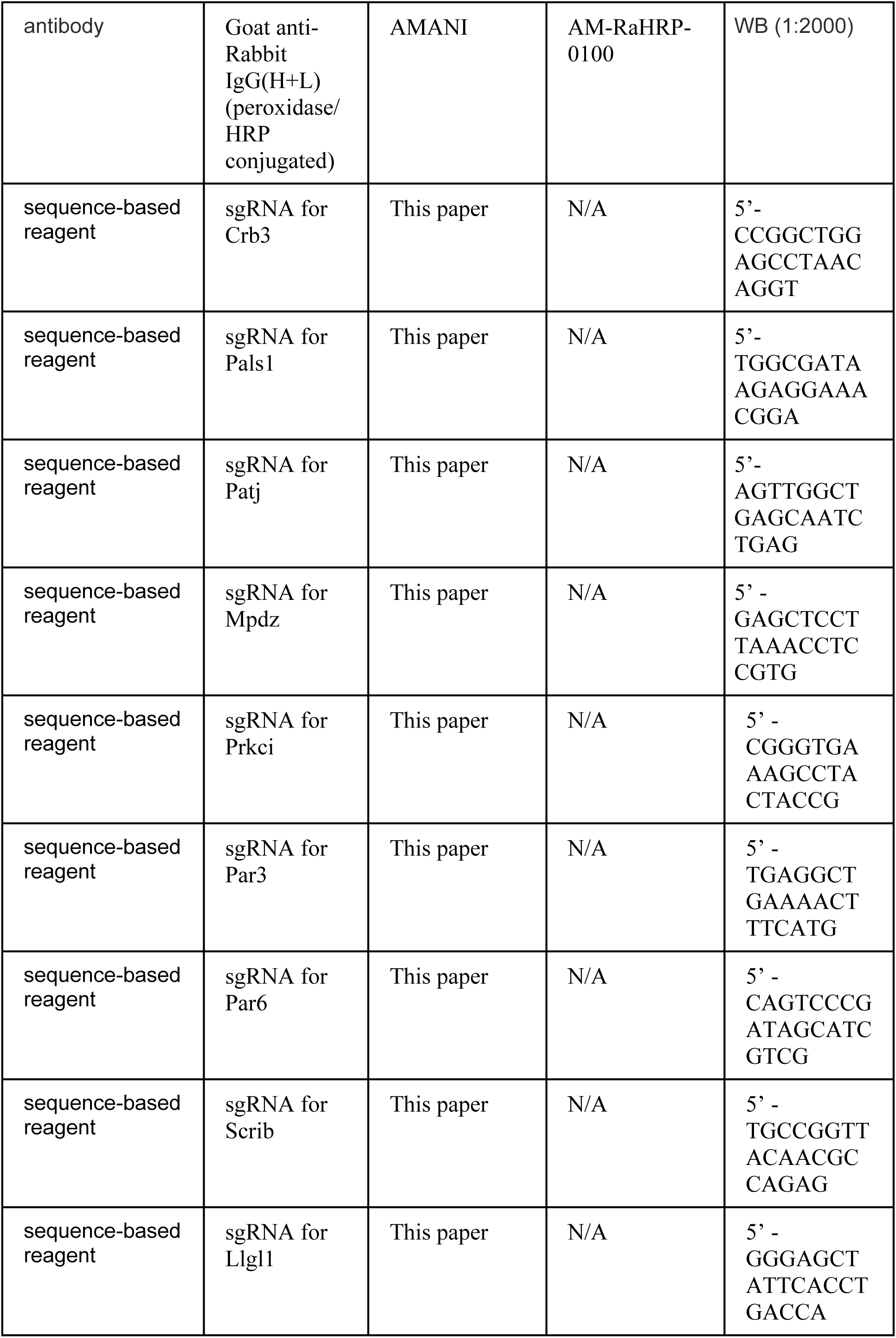
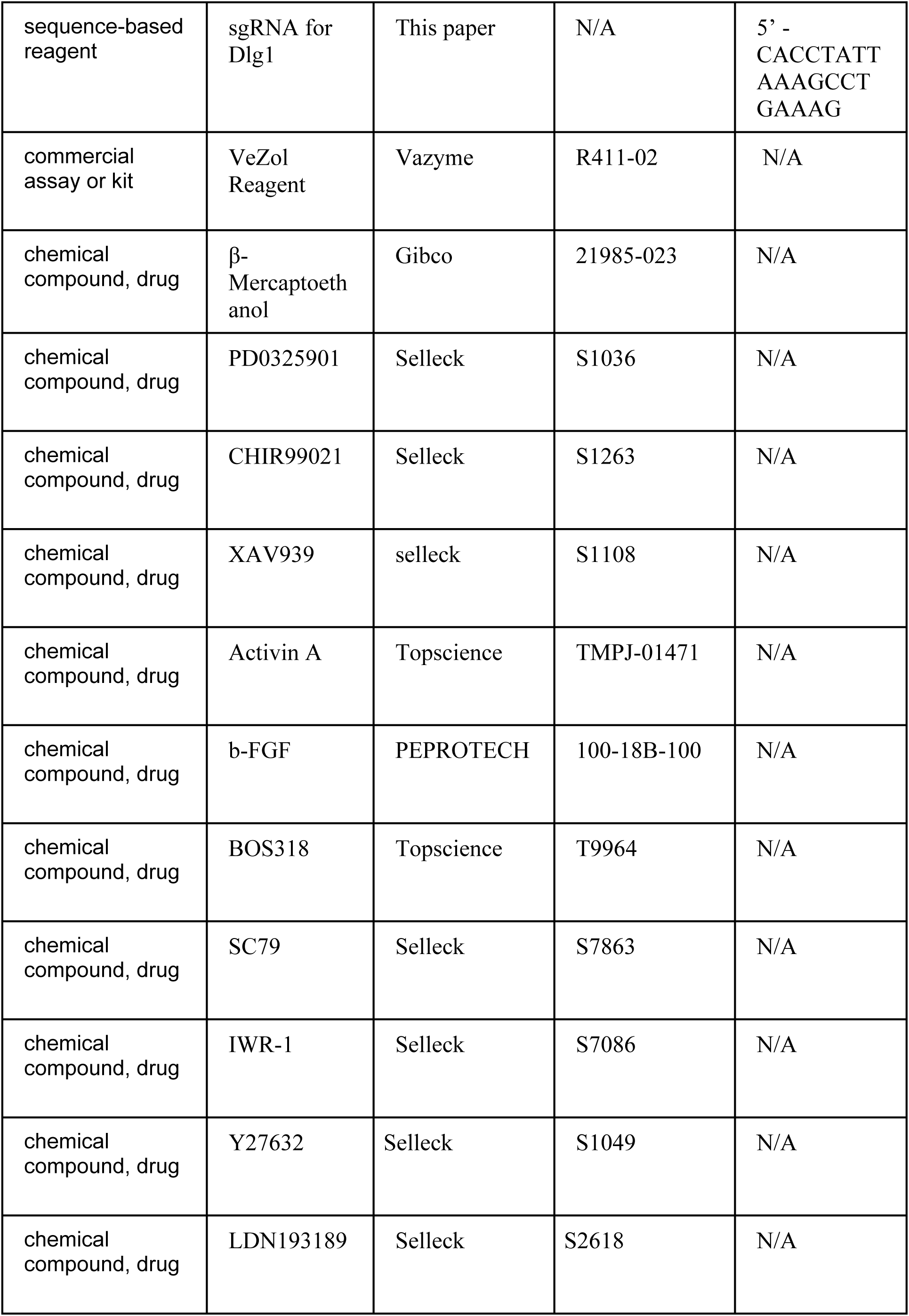
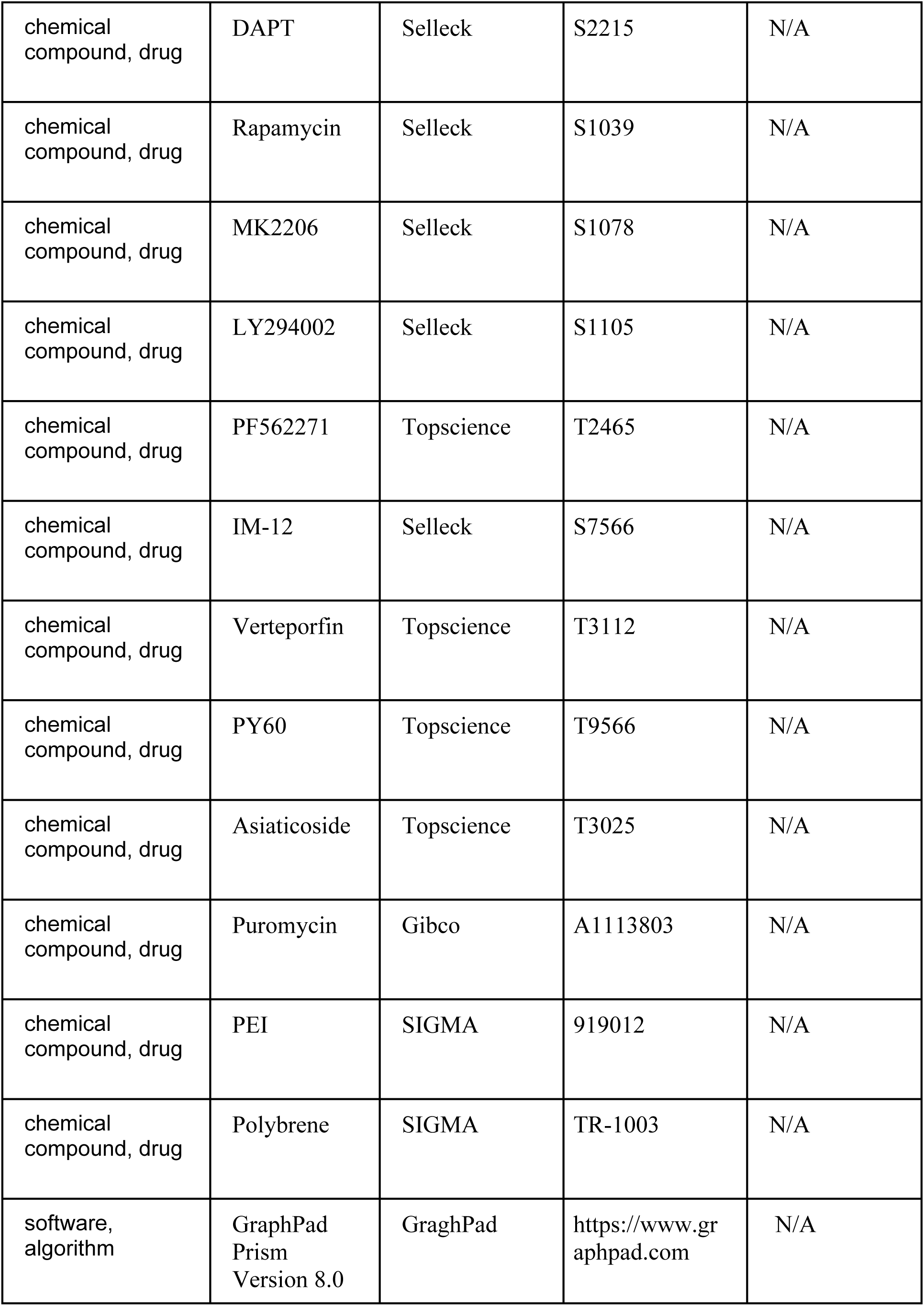
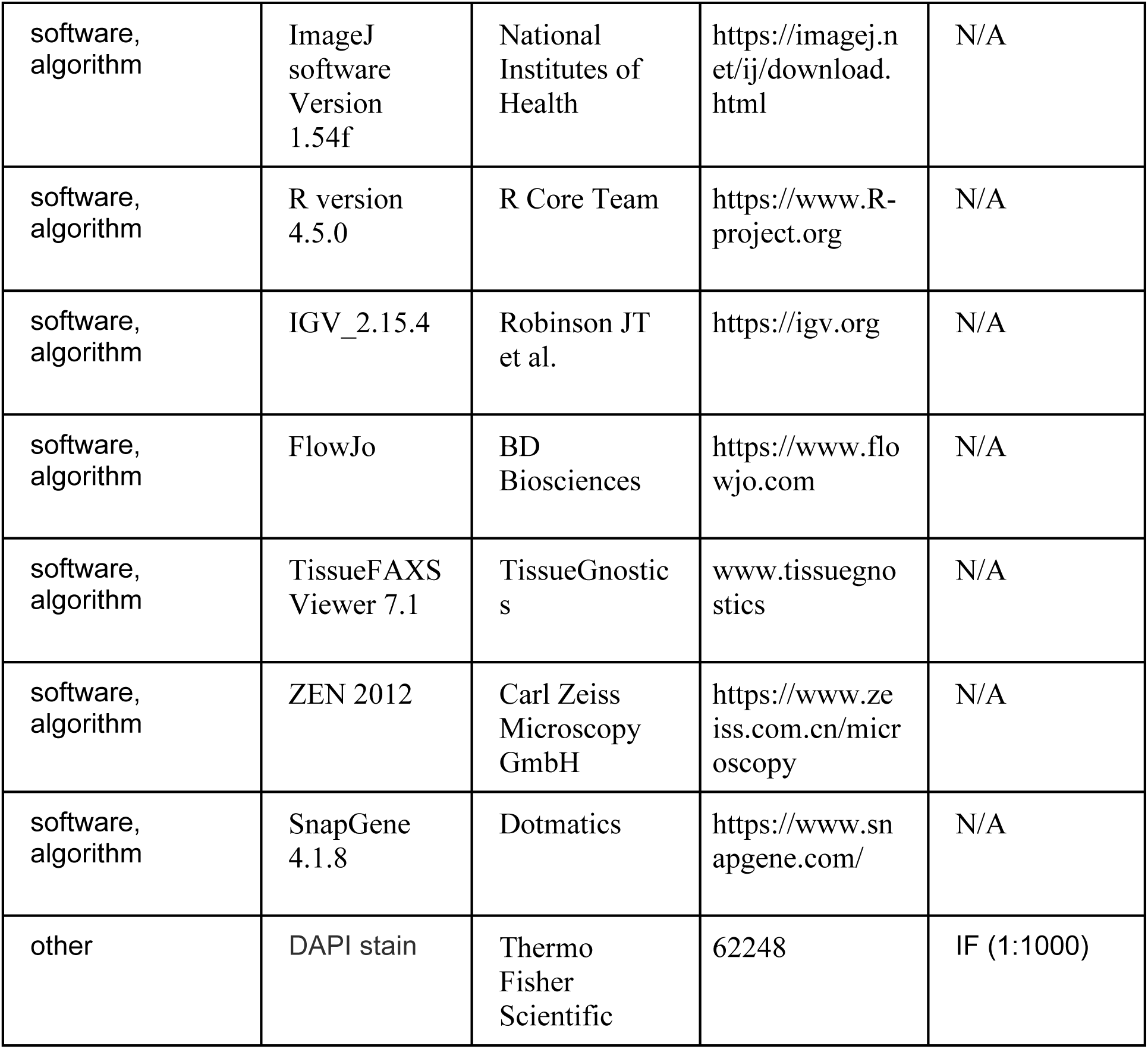
Key resources table.

## Supplementary figures

**Supplemental Figure 1.**
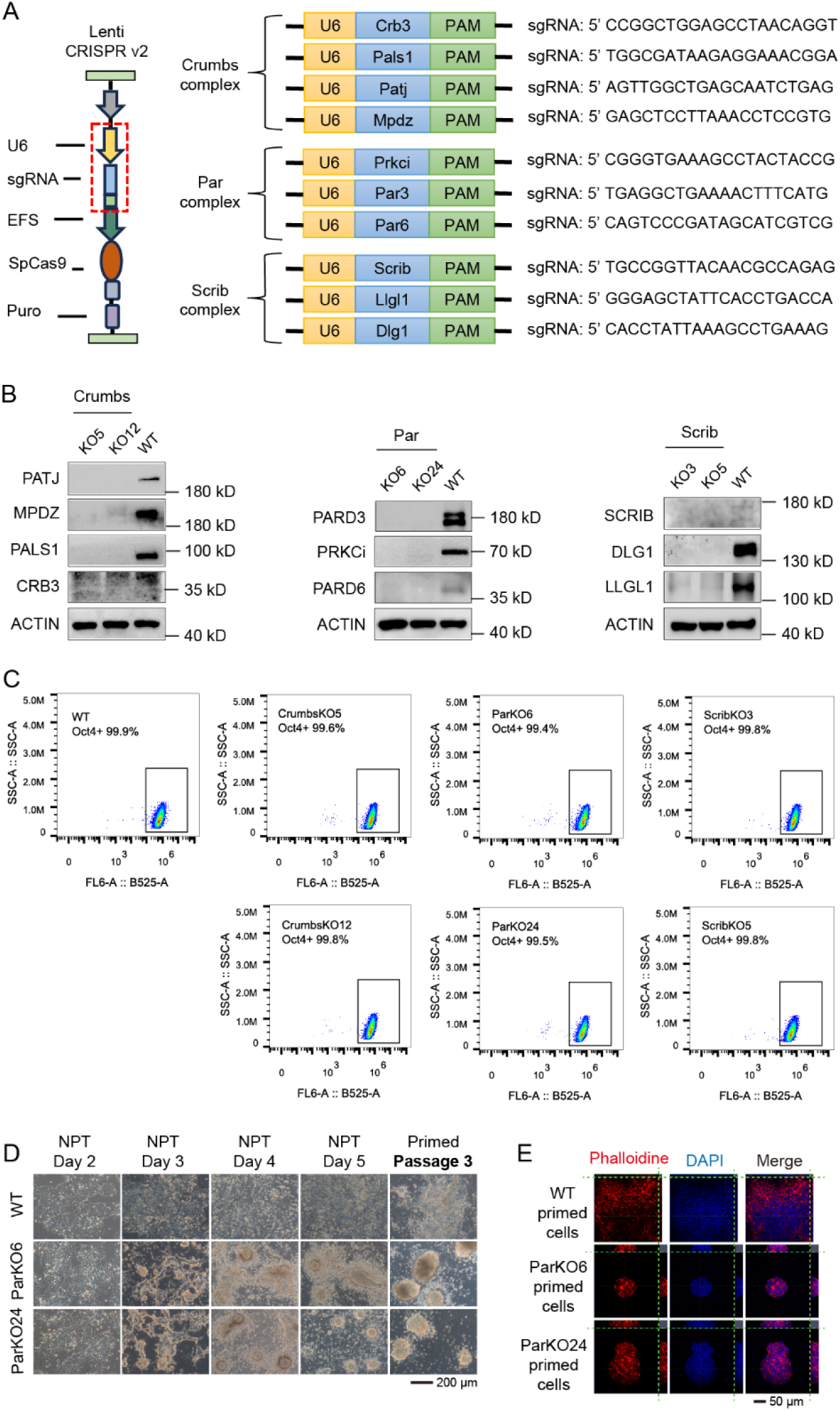
Par complex is essential for NPT, related to Figure 1. **(A)** The sgRNA design strategy for knockout of Crumbs, Par, or Scrib complex. The plasmid was constructed using the CRISPR-cas9 technology. **(B)** Knockout efficiency for Crumbs, Par, and Scrib complexes was validated by WB. **(C)** Crumbs, Par, or Scrib complex KO did not affect ESCs pluripotency, the proportion of Oct4-GFP^+^ cells was not significantly different from that of the WT. **(D)** Par complex KO caused the cells to exhibit a dome-shaped colony morphology, and this defect can be stably maintained at least three passages of culture. **(E)** WT primed cells presented flat monolayer clusters, while Par KO primed cells exhibited dome colonies, as visualized by Phalloidin staining and imaged by LSM800 confocal microscopy with Z-stack acquisition. Experiments were repeated at least three times unless otherwise stated.

**Supplemental Figure 2.**
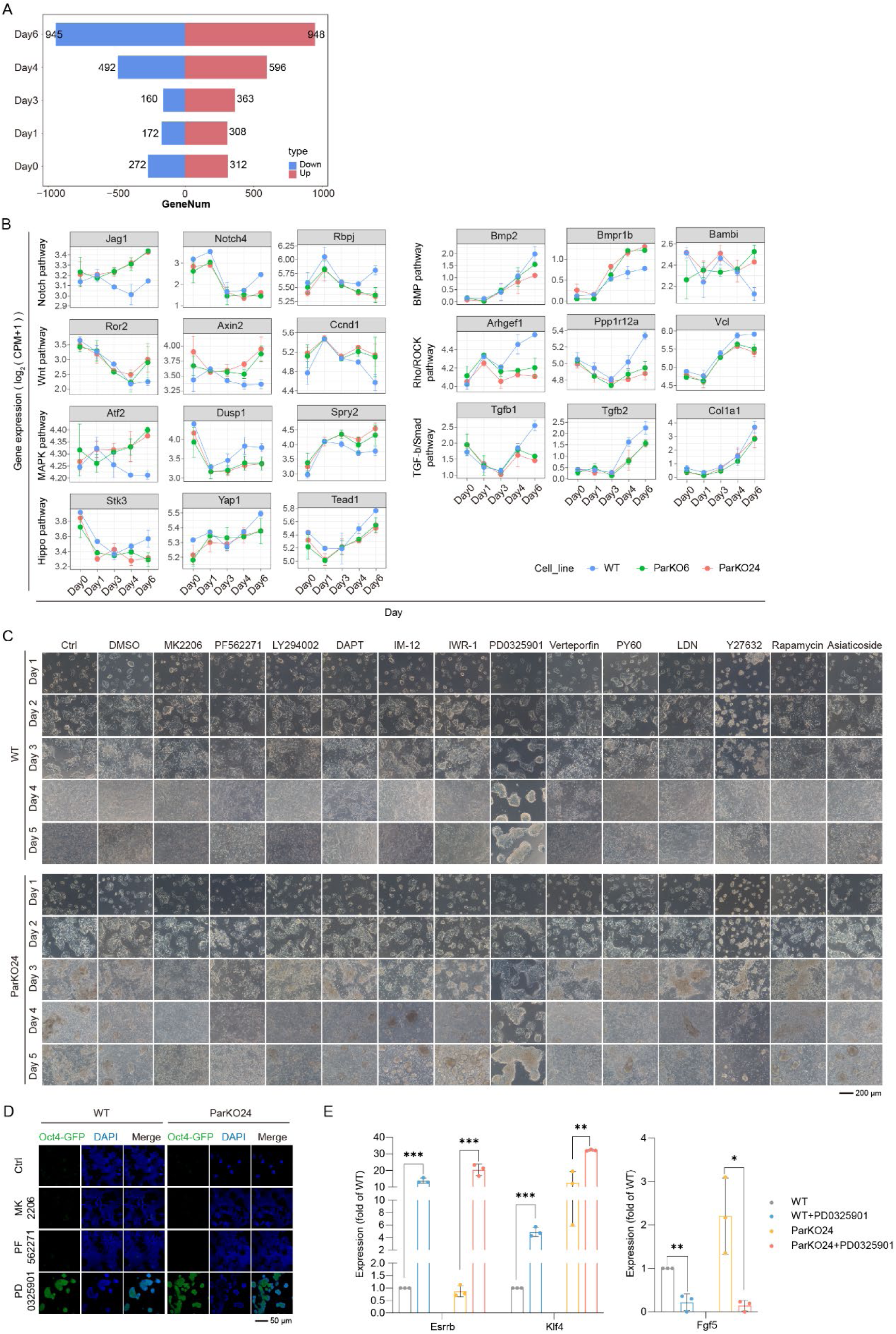
AKT and FAK are key signal pathways that regulate cell morphology during NPT, related to Figure 2. **(A)** The DEGs between Par KO and WT cells at different time points during NPT. The number of DEGs was the highest on Day 6. **(B)** Expression of key genes of the enriched pathways in **(Figure 2D)** was analyzed. **(C)** Cells were treated with agonists or inhibitors targeting the enriched pathways during NPT, followed by morphological analysis of dome colonies. **(D)** Par KO cells were treated with MK2206, PF562271, or PD0325901 during NPT and assessed Oct4-GFP^+^ expression. **(E)** The expression of ESC markers (such as *Esrrb* and *Klf4*) or primed markers (such as *Fgf5*) in cells treated with PD0325901 was analyzed during NPT. Experiments were repeated at least three times unless otherwise stated. Error bars represent S.D. Unpaired t test analysis was performed in **(E)**.

**Supplemental Figure 3.**
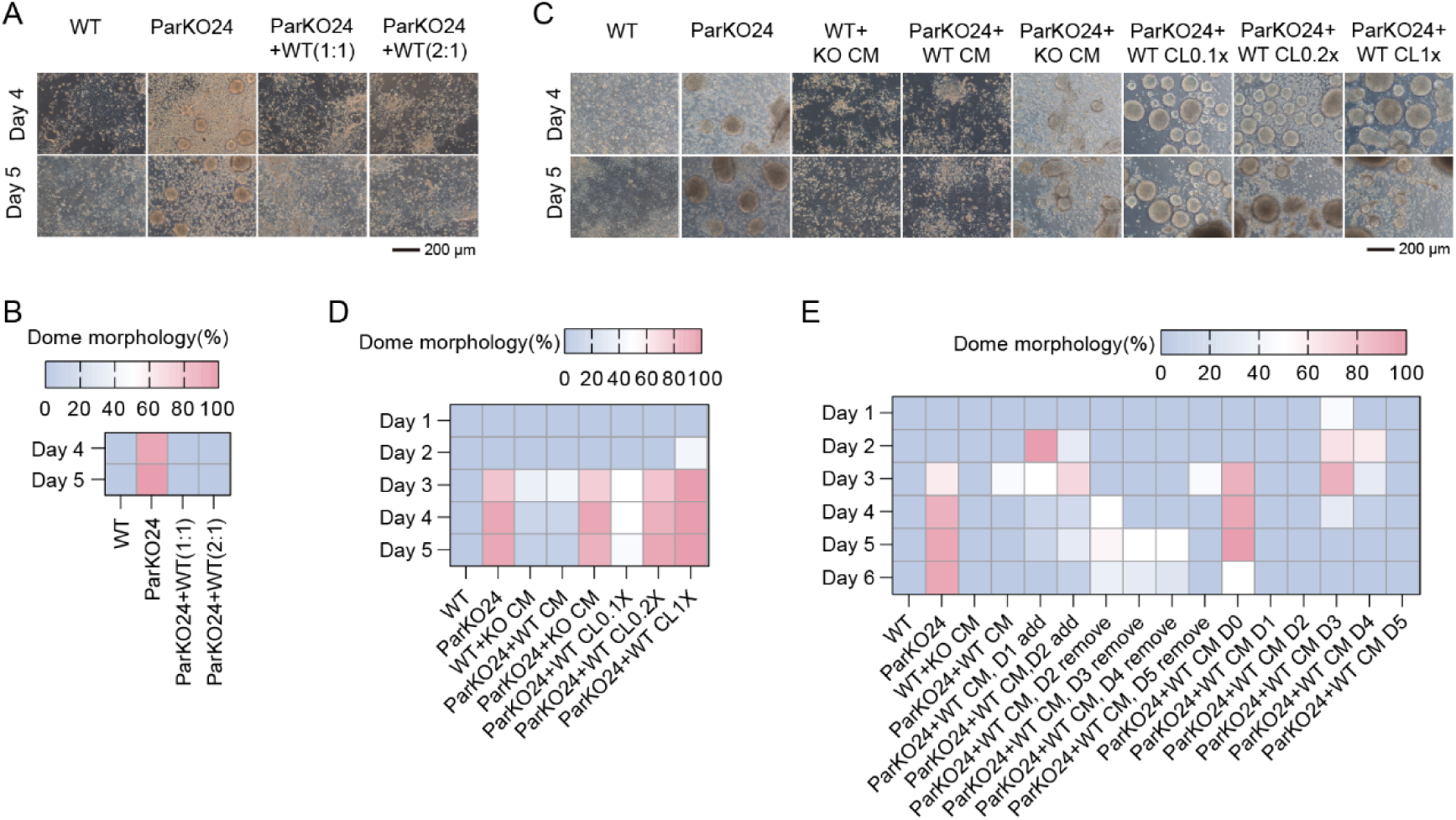
WT conditioned medium rescues the defect induced by Par KO, related to Figure 3. **(A-B)** NPT in co-culture system was carried out by mixing Par KO ESCs and WT ESCs at ratio of 1:1 or 2:1. Cell morphology was analyzed at the end of NPT. **(C-D)** Par KO ESCs were treated with WT CM, KO CM, or WT CL during NPT, and the cell morphology was analyzed. **(E)** The rescue effect of WT CM on Par KO ESCs showed a time-dependent pattern. WT CM must be present during the period from Day 2 to Day 4 of NPT to exert the rescue effect. While any collection of WT CM taken on Days 1 to Day 5 during NPT has a rescue effect. Experiments were repeated at least three times unless otherwise stated.

**Supplemental Figure 4.**
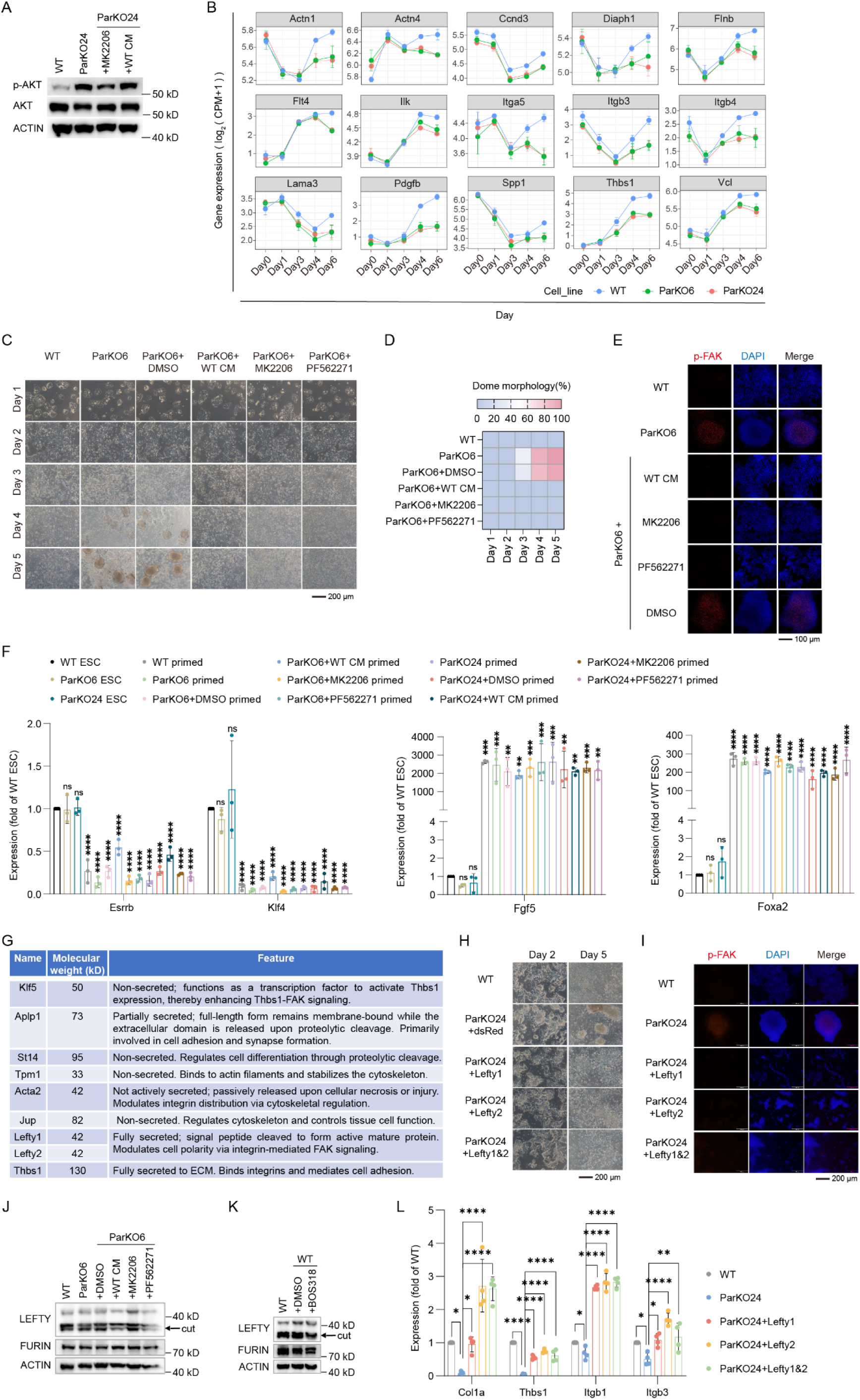
Par complex mediates cell morphology through the AKT-FAK signal axis, related to Figure 4. **(A)** Par KO cells were treated with MK2206, or WT CM during NPT, the p-AKT level were investigated on day 6. **(B)** Transcriptomic analysis of FAK-related DEGs between WT and Par KO cells during NPT using RNA-seq data. **(C-D)** ParKO6 cells were treated with WT CM, MK2206, or PF562271 during NPT, and the proportion of dome-shaped colonies was quantified. All three treatments restored the flat monolayer morphology of ParKO6 primed cells. **(E)** ParKO6 cells were treated with WT CM, MK2206, or PF562271 during NPT, the p-FAK level in the cells was detected on Day 6. **(F)** Par KO cells were treated with WT CM, MK2206, or PF562271 during NPT, and the molecular signatures of the treated cells were examined at day 6, including naïve markers (Esrrb and Klf4) and primed markers (Fgf5 and Foxa2). **(G)** Based on the FAK signaling pathway, functional candidate proteins were screened and subsequently overexpressed in Par KO ESCs to validate their biological roles. **(H)** Overexpression of LEFTY resulted in flat monolayer clusters of Par KO cells resembling that of the WT cells during NPT. **(I)** Overexpression of LEFTY in Par KO cells reduced the p-FAK level during NPT. **(J)** ParKO6 cells were treated with WT CM, MK2206, or PF562271 respectively, and the protein level of the LEFTY and FURIN was detected. The position indicated by the arrow represents the LEFTY cut by FURIN. **(K)** WT cells were treated with BOS318, the protein level of the LEFTY and FURIN was detected. **(L)** Overexpression of LEFTY in Par KO cells, followed by qPCR analysis of mRNA levels of ECM-related genes (Col1a, Thbs1) and integrin-related genes (Itgb1, Itgb3) during NPT. Experiments were repeated at least three times unless otherwise stated. Error bars represent S.D. One-way ANOVA was performed in **(F)** and **(L)**.

**Supplemental Figure 5.**
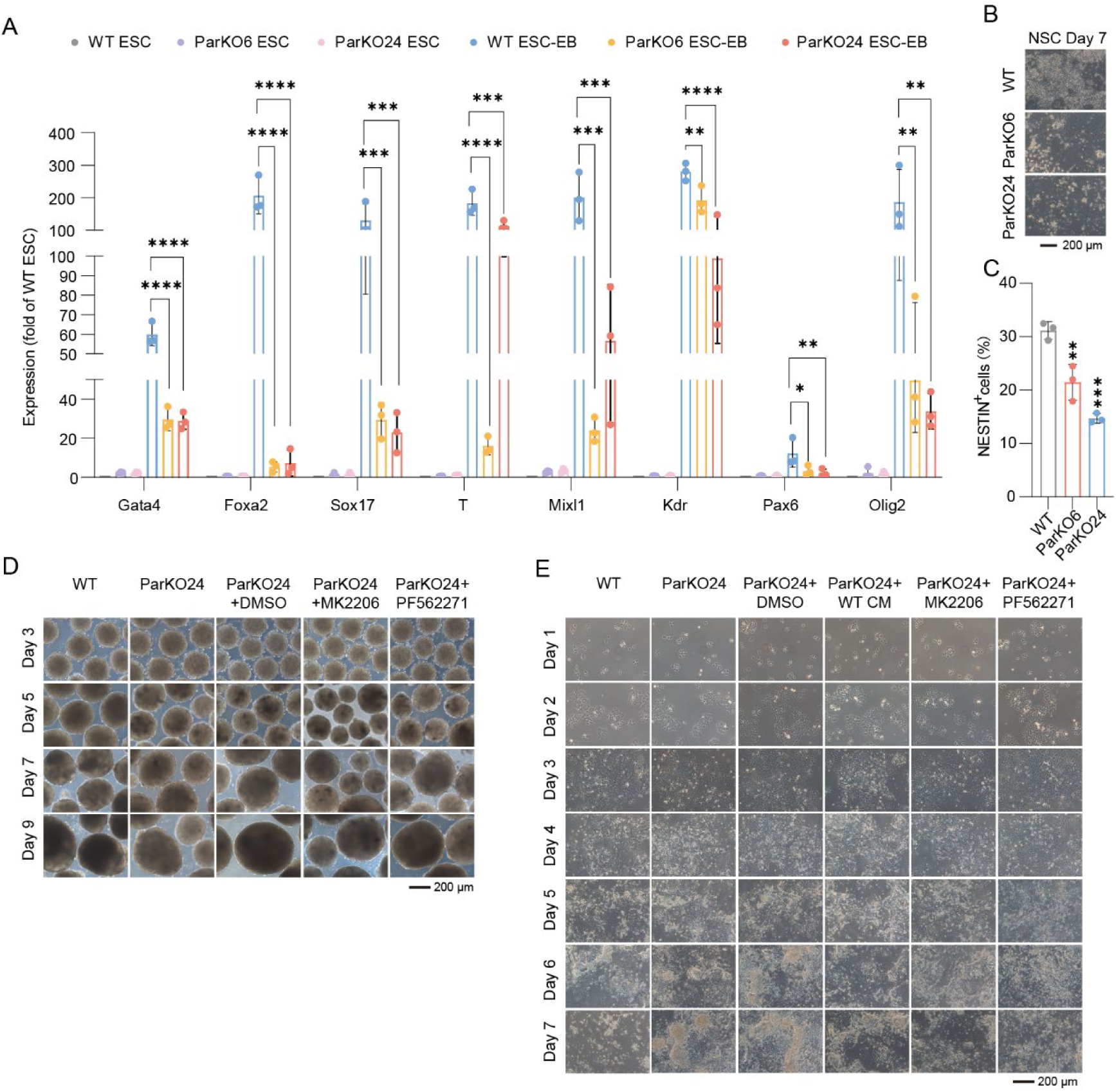
Par KO impairs lineage differentiation via the AKT–FAK signaling axis, related to Figure 5. **(A)** During EB differentiation from ESCs, expression of three germ layer markers was detected. **(B-C)** ESCs were induced to differentiate into NSCs. The cell morphology was analyzed, and the expression of NESTIN, a marker for NSCs, was detected. **(D)** Par KO ESCs were treated with MK2206 or PF562271 during the EB differentiation. The EB differentiation was performed using a suspension culture method. **(E)** Par KO ESCs were treated with WT CM、MK2206 or PF562271 during the NSC differentiation. Experiments were repeated at least three times unless otherwise stated. Error bars represent S.D. One-way ANOVA was performed in **(A)** and **(C)**.

**Supplemental Figure 6.**
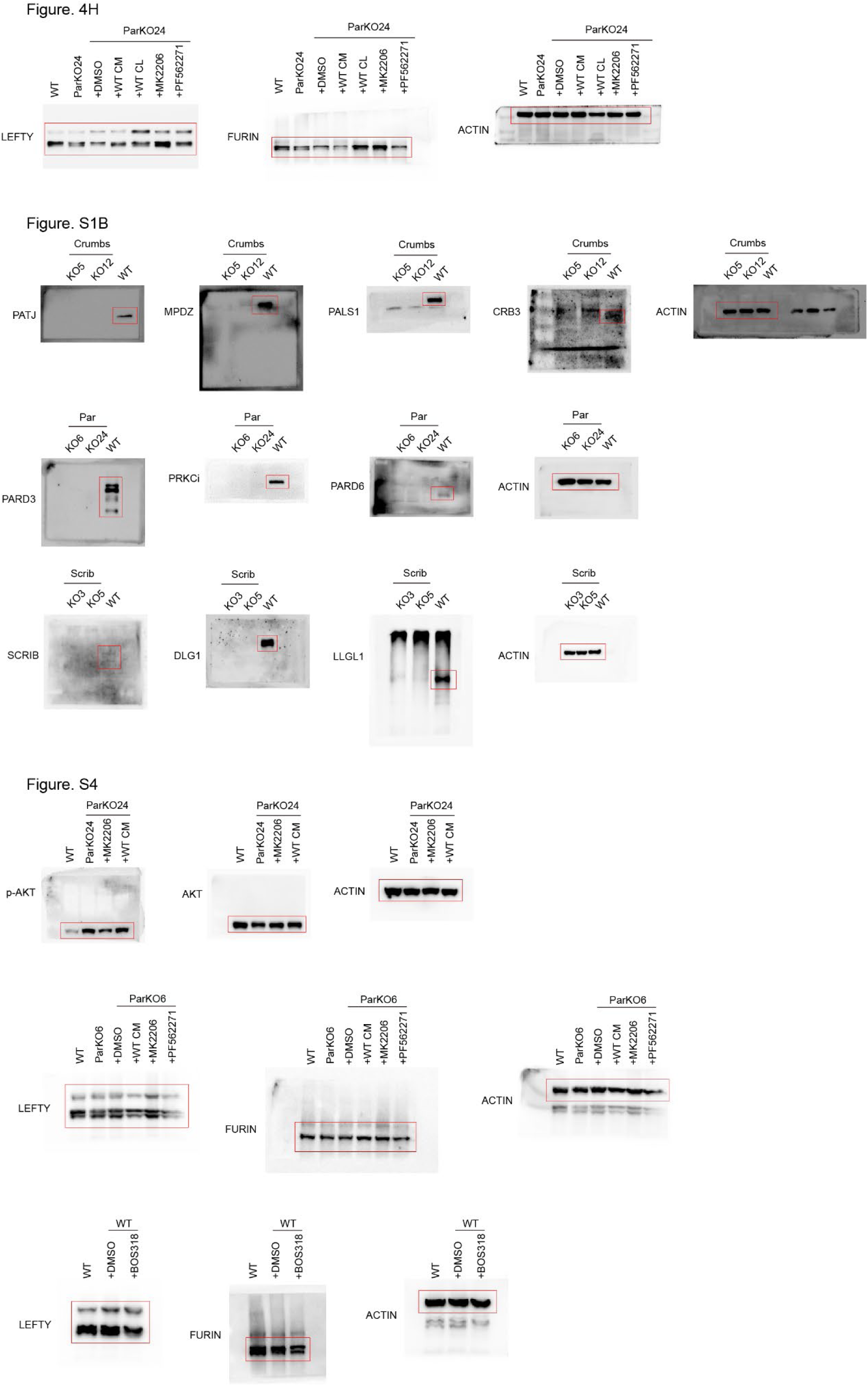
Uncropped gels, related to Figure 4, Figures S1 and S4.

## Acknowledgements

The projects were supported by National Key R&D Program of China (2024YFA1108201 and 2024YFA1802303), the Strategic Priority Research Program of Chinese Academy of Sciences (XDB1030303), Guangdong Basic and Applied Basic Research Foundation (2024B1515040020 and 2024A1515010609), Science and Technology Program of Guangzhou (202201011654), Science and Technology Planning Project of Guangdong Province (2023B1212060050), Guangdong Special Support Program (2023TX07A051), Postdoctoral Research Project Funding of He Meiai (B202500701), the Youth Innovation Promotion Association of the Chinese Academy of Sciences (2022362), the Tertiary Education Scientific research project of Guangzhou Municipal Education Bureau (2024312185), Human Cell Lineage and Atlas Facilities (DSS010403-01), and Research Funds from Health @InnoHK Program launched by Innovation Technology Commission of the Hong Kong SAR, P. R. China.

## Additional information

### Lead contact

Further information and requests for resources and reagents should be directed to and will be fulfilled by the lead contact, Hui Zheng.

### Materials availability

All materials, reagents, and transgenic lines generated in this article are available upon request to the lead contact.

### Data and code availability

All raw data generated in this study have been deposited in the China National Center for Bioinformation/National Genomics Data Center under GSA: CRA032782 and OMIX: OMIX012794. All data are available in the main text or the supplementary materials. The authors are willing to distribute all materials, datasets, and protocols described in this manuscript. Dr. Hui Zheng assumes responsibility for responding to requests and providing information regarding reagents and resource sharing. Further information and requests for resources and reagents should be directed to and will be fulfilled by Hui Zheng.

This paper does not report original code.

All information needed to reanalyze the data reported in this paper will be shared by the lead contact upon request.

### Author contributions

Conceptualization, H.Z., L.N.L., D.J.Q.; Data Curation and Formal Analysis: M.A.H., Y.L.W., H.S., L.N.L.; Investigation: M.A.H., Y.Y.C., J.C.H., Y.H.W., Q.W.R., L.N.D., L.L., Q.Q.Z., T.Y.Z., X.Y.H., Q.G.; Supervision: H.Z., L.N.L., D.J.Q., L.G., C.P.L., S.Y.C.; Writing—Original Draft: M.A.H., L.N.L.; Writing—Review & Editing: M.A.H., L.N.L., H.Z.

